# Regional Cell Atlas of Human Intestine Shapes Distinct Immune Surveillance

**DOI:** 10.1101/2021.09.13.459999

**Authors:** Yue Wang, Yanbo Yu, Lixiang Li, Mengqi Zheng, Jiawei Zhou, Haifan Gong, Bingcheng Feng, Junyan Qu, Zhen Li, Rui Ji, Ming Lu, Xiaoyun Yang, Xiuli Zuo, Shiyang Li, Yanqing Li

**Author notes:** Corresponding author. (XZ); (SL); (YL). These authors contributed equally to this work (Yue Wang, Yanbo Yu, Lixiang Li, Mengqi Zheng).

## Abstract

Regional intestinal immune surveillance remains obscure. In this study, we integrated single-cell RNA sequencing and spatial transcriptomics to create a regional atlas of fetal and adult intestines, consisting of 59 cell subsets, of which eight new subsets and ILCs transition states were identified. Results revealed that microenvironment determines in-situ cell differentiation and shapes the regional molecular characteristics, allowing different intestinal segments with diverse functions. We characterized the regional expression of mucins, immunoglobulins, and antimicrobial peptides (AMPs) and their shift during development and in inflammatory bowel disease. Notably, α-defensins expressed most abundantly in small intestinal *LGR5*^+^ stem cells, rather than in Paneth cells, and down-regulated as cell maturing. Common upstream transcription factors controlled the AMPs expression, illuminating the concurrent change of AMPs during epithelial differentiation, and the spatial co-expression patterns. We demonstrated the correspondence of cell focus of risk genes to diseases’ location susceptibility and identified distinct cell-cell crosstalk and spatial heterogeneity of immune cell homing in different gut segments. Overall, a cross-spatiotemporal approach to transcriptomes at single-cell resolution revealed that the regional milieu of the human intestine determined cellular and molecular cues of immune surveillance, dictating gut homeostasis and disease.

## Introduction

Intestinal immune surveillance is essential for coordinating nutrition and immunity between the body and the environment since it mediates microbial tolerance and pathogen defense^1–5^. Immune-surveillance disorders are linked to diseases with unique regional susceptibility, such as inflammatory bowel disease (IBD) and gastrointestinal tumors^6–9^.

Recent single-cell RNA sequencing (scRNA-seq) studies have provided new insights into the dysfunction of mucus barrier and immune cells during diseases with regional susceptibility^10–15^. Nevertheless, the regional patterns of intestinal immune surveillance for microorganisms and the cellular, molecular, and intercellular mechanisms that support the surveillance remain unclear. In addition, the shifts in immune surveillance during fetal development and in diseases need to uncovered.

Mucus layer, epithelium, and immune cells constitute the hierarchical immune surveillance in human intestinal tract^16–18^. In this study, we aimed to integrate scRNA-seq and spatial transcriptome (ST) to create a comprehensive intestinal cell atlas of fetus, healthy adults, and patients with IBD, to identify the molecular mechanisms underlying the shaping of epithelial cell differentiation and function. Our data revealed the contribution of regional milieu in the course of intestinal diseases by delineating the regional heterogeneity of immune surveillance, and its shift during fetal development and in IBD.

## Results

### High-resolution cell atlas of the healthy human intestine

A total of 14 biopsy samples, representing duodenum, jejunum, and ileum, were collected and segregated into epithelium or lamina propria, and single-cell transcriptome sequencing, based on 10x Chromium, was performed (Extended Data Fig. 1a), generating high-quality transcriptomes from 53,749 cells. The new generated profiles coincided with the published data of 54 human samples (37,821 cells) from healthy ileum, colon, and rectum^10,19^, forming a comprehensive cell atlas covering epithelial, immune, and stromal compartments (Extended Data Fig. 1b).

According to compartment-specific markers, unsupervised clustering preliminarily divided the cells into six compartments, namely epithelium, T/innate lymphoid cells (ILCs), B cells, stromal cells, mononuclear phagocytes (MNPs), and mast cell compartments (Fig. 1a, b, Extended Data Figs. 1c, 2a). Based on iterative clustering, 59 subsets were identified with unique gene expression and segment distribution (Fig. 1c, f, Extended Data Figs. 1d, 2b-c, 3a-c), of which eight were newly defined (Fig. 1e).

**Fig. 1:**
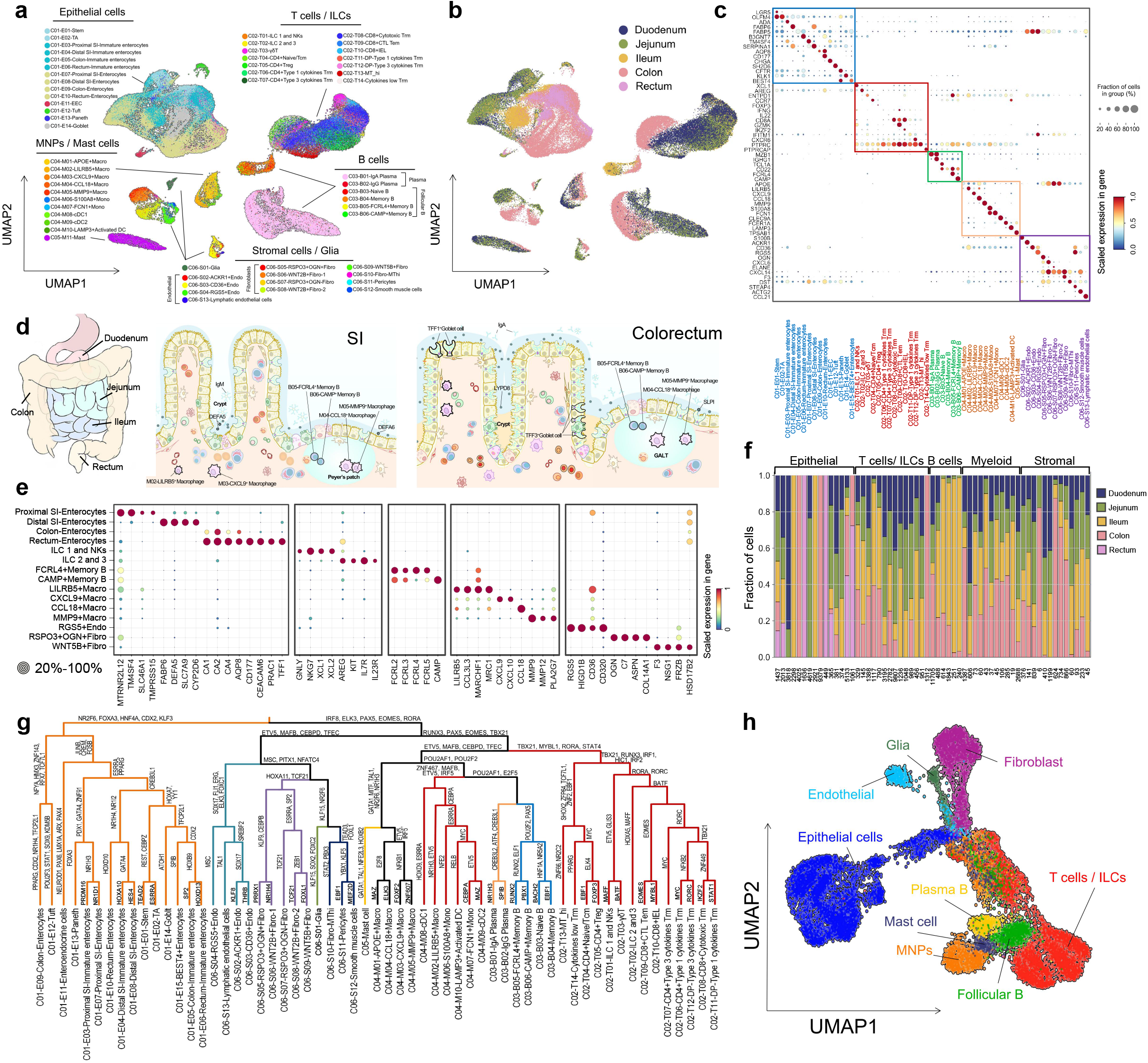
Single-cell atlas of the healthy human intestine. **a**, UMAP embedding of cells from 59 different cell subsets. **b**, UMAP embedding overlay showing the location distribution across all cell subsets. **c**, Subset-specific markers. Shown is the scaled expression of marker genes (rows) across cell types (column) (see also in Extended Data Figs. 2, 3). **d**, Diagrams showing localization and morphology of newly identified cell subsets, and differentially expressed immunoglobulins and antimicrobial peptides across different intestinal regions. **e**, Dot plot showing markers of 8 newly identified cell subsets, ILCs transition state, and enterocytes from different intestinal regions. **f**, Cell subset distributions. Shown are fractions of cells (y-axis) in each cell subset (bars) that are derived from indicated samples. Bottom: total cell counts in the subset. **g**, Hierarchical clustering of all cell types based on TF module scores. Branches are colored by compartment. TFs at each branching point of the dendrogram are representative for subjacent groups of regulons (see also in Extended Data Fig. 5d and Supplementary Table 2). **h**, UMAP embedding based on the TF module scores shows the intestinal cells into identical but consecutive cell lineages (see also in Extended Data Fig. 5a, b).

### New intestinal cell subsets and states

M02-*LILRB5*^+^ macrophages (*LILRB5*^+^*CCL3L3*^+^*MRC1*^+^*CD163*^+^) were enriched in the small intestine (SI), characterized by specific chemokines and receptors (*CCL3L3, CCL4L2, CCR1, CX3CR1*, and *C5AR1*) (Fig. 1e, f, Extended Data Fig. 3c), and were scattered in lamina propria, as revealed by spatial transcriptome analysis (Extended Data Fig. 1e). Intestinal *LILRB5*^+^ macrophages, capable of putative self-recruitment through the *CX3CR1-CCL3L3/CCL4L2* axis, might sense bacterial metabolites and limit inflammation via CX3CR1 and *CD163*, respectively^20,21^. M03-*CXCL9*^+^ macrophages (*CXCL9*^+^ *CXCL10*^+^), populated in the terminal ileum and colon (Fig. 1e, f, Extended Data Fig. 3c), might correlate with intestinal inflammation by interacting with cDC1, Th1-like Trm, and DP + Th1-like cells through *CXCL9/CXCL10-CXCR3*. In contrast to M02 and M03, M04-*CCL18*^+^ (*CCL18*^+^) and M05-*MMP9*^+^ (*MMP9*^+^*MMP12*^+^*PLA2G7*^+^) macrophages localized in gut-associated lymphoid tissue (GALT) (Fig. 1e, Extended Data Fig. 1e). The transmission and differentiation of M05 might be controlled by matrix metalloproteinases 9 and 12^22,23^.

B05-*FCRL4*^+^ and B06-*CAMP*^+^ memory B cells mainly located in the GALT of the ileum (Fig. 1e, f, Extended Data Figs. 1e, 3b). Highly expressed chemokine receptors (*CCR1*, *CCR2, CCR6*, and *FCGR2A*) in B05 might enhance their homing during inflammation (Extended Data Fig. 3b) ^24^, whereas LL-37, an AMP encoded by *CAMP*, might cause B06 to exhibit anti-bacterial function^25,26^.

A rare but unique subset, S06-*RGS5*^+^ endothelial cells, presented a functional state of hypoxia-induced apoptosis in endothelial lineage, as suggested by the up-regulation of *RGS5, SOD3*, and *GPX3* (Fig. 1e, Extended Data Fig. 2c)^27,28^. SI-populating S05-*RSPO3*^+^*OGN*^+^ fibroblasts expressed similar gut homeostasis-supporting genes (Fig. 1e, f, Extended Data Fig. 2c) with S07-*RSPO3*^+^*OGN*^-^ fibroblasts, which had been identified previously in colon^10^. However, S05-*RSPO3*^+^ fibroblasts up-regulated neutrophils- and T cell-homing chemokines, *CXCL2, 3, 6*, and *16* while down-regulating the plasma cell-recruiting chemokine *CCL7*. Notably, S05 expressed higher *C3* and *C7* than colon S07.

Lineage-specific genes of NK, ILC1, ILC2, and ILC3 enriched in two independent cell clusters, namely NK-ILC1 and ILC2-3 (Extended Data Fig. 4a-d). ILC2-3 subset co-expressed ILC2-related genes, *AREG* and *GFI1*, as well as ILC3 markers, *RORC* and *IL22* (Fig. 1e, Extended Data Fig. 4a-d). Failed attempt to regroup NK-ILCs, along with continuous state reflected by the pseudo-time analysis (Extended Data Fig. 4b-d), suggested the existence of a transitional state of NK-ILC compartment and was further confirmed by fluorescence-activated cell sorting (FACS) (Extended Data Fig. 4e-g).

### Fate decision tree of intestinal cells

A gene regulatory network (GRN), composed of 543 activated regulator modules, was constructed^29,30^. We identified conical transcription factors (TFs) of known cell types, such as *ATOH1* for secretory enterocytes and *RORC* for *CD4*^+^ and DP-type 3 cytokine T cells, and TFs of newly identified cell types, such as *POU2F2* for *FCRL4*^+^ memory B cells and *ZNF732* for *LILRB5*^+^ macrophages (Extended Data Fig. 5d). Hierarchical clustering and differential analysis found regulon switches that determined the differentiation of cell subsets (Fig. 1g, Extended Data Fig. 5d). For example, *SOX17* and *TAL1* controlled the differentiation of lymphatic and vascular endothelial cells, respectively, followed by *MSC* activation to promote *RGS5*^+^ endothelial cell generation. In contrast to the whole transcriptome, regulon activity grouped the intestinal cells into identical but consecutive cell lineages (Fig. 1a, h).

### Shaping of regional epithelial cell differentiation and molecular characteristics by the microenvironment

The anatomical position of intestine deeply affected the differentiation of regional epithelial subsets based on 41 unique GRNs and fate decision trees (Fig. 2a), i.e., *GATA4, GATA5*, and *PDX1* for SI, and *TFCP21*, *HOXB9*, and *HOXB13* for colon stem cells, transit-amplifying (TA) cells, and immature enterocytes, respectively. However, the sensitivity of different epithelial subset GRNs to the microenvironment was variable. Absorptive enterocytes, goblet cells, and Paneth/*BEST4*^+^ enterocytes showed significant GRN heterogeneity across the intestine, although not so in tuft cell and enteroendocrine cells (EEC) GRNs. Bile acid receptors *NR1H4* and *NR1H3*^31,32^ were highly expressed in small intestinal stem cells, TA cells, and immature enterocytes, and less so in mature goblet cells, enterocytes, and Paneth cells, indicating the involvement of environmental signals in epithelial cell differentiation since bile acids differ regionally^33^.

**Fig. 2:**
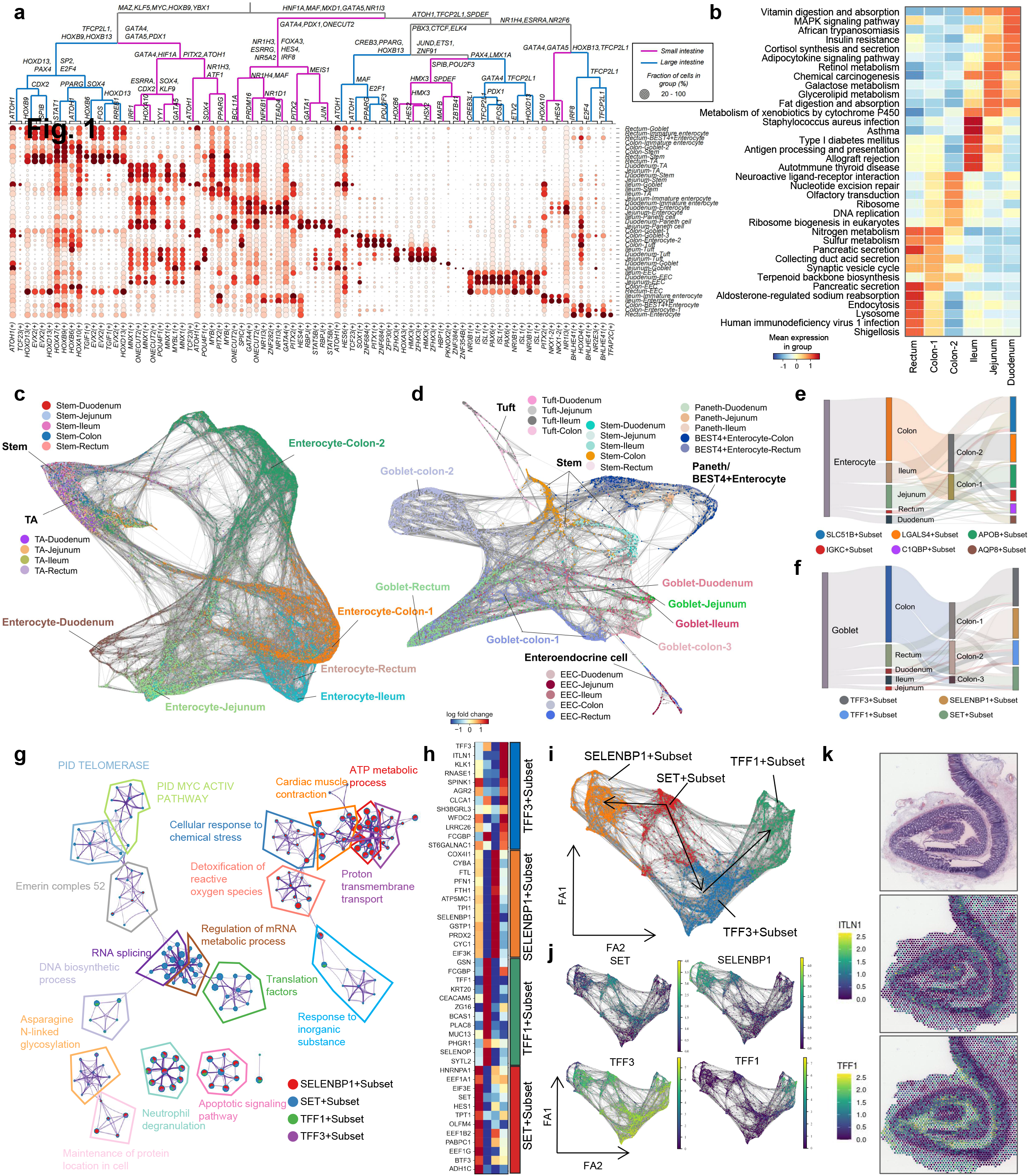
The microenvironment determines in site epithelial cell differentiation and shapes regional molecular characteristics. **a**, A dendrogram of regulons for 41 regional epithelial cell subsets. TFs at each branching point of the dendrogram are representative of subjacent groups of regulons. Branches are colored by location (see also in Supplementary Table 3). **b**, Regional function shifts in enterocytes across different intestinal regions. Shown are scaled scores of KEGG pathways (rows) (see also in Extended Data Fig. 6c-e). **c**, **d**, Trajectory and partition- based graph abstraction of absorptive (c) and secretory (d) epithelial cell lineages. **e**, **f**, Sankey diagrams depicting the mapping of enterocytes (e) and goblet cells (f) from different intestinal regions to their functional subsets. **g**, **h**, Functional (g) and molecular (h) characteristics of goblet functional subsets. **i**, **j**, Trajectory and partition-based graph abstraction highlighting the differentiation of goblet functional subsets (i) and their marker genes (j). **k**, Expression of *ITLN1* and *TFF1* in ST adult colon slides and H&E image showing the positioning on the crypt-villus axis of *TFF3*^+^ goblet subset and *TFF1*^+^ goblet subset.

Consistently, the pseudo-time analysis revealed that regional heterogeneity of enterocytes, goblets, and *Paneth/BEST4*^+^ enterocytes was higher than that of tuft cells and EECs, increasing during cell differentiation (Fig. 2c, d). Enterocytes, goblets, and Paneth/*BEST4*^+^ enterocytes showed distinct functional enrichment, according to the local environment of proximal SI, ileum, and colorectum (Fig. 2b, Extended Data Figs. 6c-e, 7e-g, 8d-f). Enterocytes in fat-rich proximal SI showed up-regulation of genes associated with fat-soluble vitamins’ digestion and absorption pathways, such as vitamin A (Fig. 2b). In contrast to their enrichment in digestive, absorption, and metabolic pathways of the proximal SI, ileal enterocytes were enriched in a variety of immune-related processes (Fig. 2b, Extended Data Fig. 6c), in line with the excessive lymphoid structures in the ileum^34^. It is noteworthy that enterocytes converged to goblet cells in mucus biosynthesis and secretion toward the end of the digestive tract, based on the up-regulation of various mucus-related genes, such as *SPINK4, CLCA1*, and *FCGBP*, as well as mucus-related metabolic pathways (Extended Data Fig. 6b, d). Enriched goblet cells and goblet-like enterocytes in the distal intestine might constitute a strengthened mucus barrier (Extended Data Figs. 2b, 6b, d), adapted to the microbiota-shaped environment.

### Functional epithelial subset differentiation

Next, we investigated the functional subsets of intestinal epithelium that are influenced by the microenvironment. Mature enterocytes, goblets, and Paneth/*BEST4*^+^ enterocytes were regrouped into 6, 4, and 6 functional subsets, respectively, with a pan-intestine distribution but extreme regional heterogeneity (Fig. 2e, f, Extended Data Figs. 6a, 7a-d, 8a-c). Take goblet cell as an example, rectum-enriched *SET*^+^ goblet subset was premature, characterized by active RNA splicing, mRNA metabolism, and growth factor response. *SELENBP1*^+^ goblet cells, mainly populated in the colon, represented higher injury or apoptosis, as revealed by enrichment of cell response to chemical stress and detoxification of reactive oxygen species (Fig. 2g, i, Extended Data Fig. 7a-d). Both *TFF3*^+^ and *TFF1*^+^ goblet cells functioned in mucus formation and microbial defense. *TFF3*^+^ goblet cells, enriched in the bottom of crypts in the colon, expressed increased levels of mucus-related (*RNASE1, AGR2, TFF3, CLCA1*, and *SPINK1*) and microbial defense-related (*ITLN1* and *WFDC2*) genes (Fig. 2h, k). However, *TFF1*^+^ goblet cells, mainly located at the top of SI crypts, up-regulated *TFF1, FCGBP, ZG16*, and *LYPD8* expression levels, which were involved in mucus layer stability and microbial defense (Fig. 2h, k). Notably, the pseudo-time analysis suggested that the *TFF1*^+^ subset was derived from the *TFF3*^+^ subset (Fig. 2i, j, Extended Data Fig. 7b-d).

### Antimicrobial peptide-mediated regional immune surveillance

Analysis of 40 gut-expressing AMPs demonstrated that the expression pattern of most AMPs varied in different intestinal regions (Fig. 3a), such as *DEFA5/6, REG3A, REG3G, ITLN2, DMBT1, GBP1, LEAP2*, and *H2BC6/7/8* in SI, and *WFDC2, SLPI, LYPD8, CCL28, ADM, DEFB1, PI3, CCL20, CXCL1, CXCL2, H2BC12/21*, and *RNASE6* in the large intestine (LI). Both SI and LI showed higher AMPs in the distal part, except for *H2BC12*. In addition, AMPs expression showed cell specificity, i.e., *HAMP* in proximal SI EECs, *TAC1* in ileal EECs, and *NPY* in proximal SI Paneth cells. Notably, region-specific AMPs were expressed in all epithelia of specific segments, though the expression levels differed (Fig. 3a). Taken together, the regional and cell type-specific AMP expression patterns seemed to shape the microbial community.

**Fig. 3:**
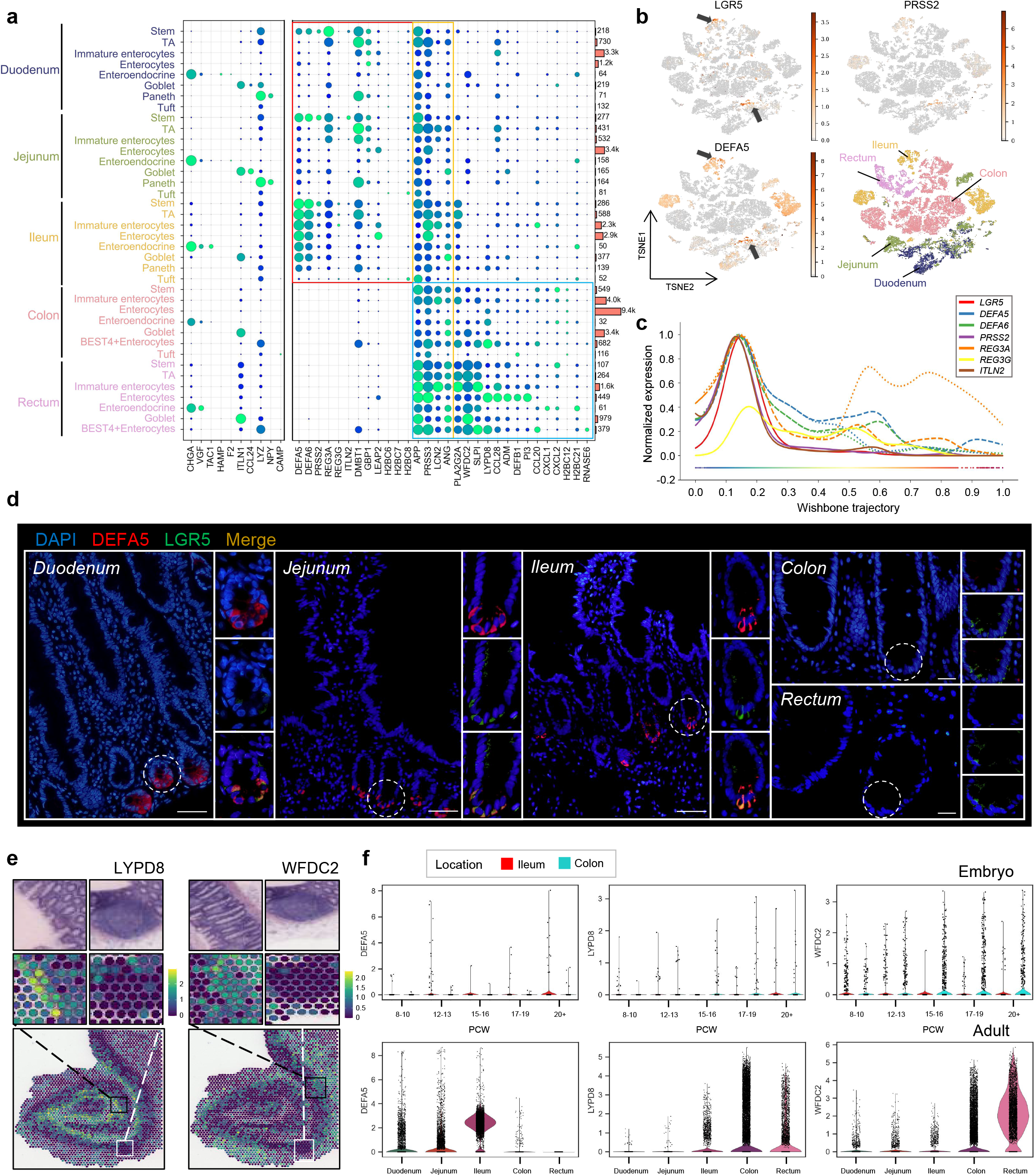
Antimicrobial peptide-mediated immune surveillance. **a**, Dot plot showing the diversity of AMPs expression pattern in different intestinal regions and cell specificity of AMPs. **b**, T-SNE embedding of epithelial cells showing the co-expression of small intestine-specific AMPs genes such as *DEFA5* and *PRSS2*, and marker gene of stem cell (i.e. *LGR5*) in duodenum, jejunum, and ileum. **c**, The wishbone trajectory dendrogram of duodenum epithelial cells highlighting the expression of small intestine-specific AMPs genes (*DEFA5*, *DEFA6*, *PRSS2*, *REG3A*, *REG3G*, *ITLN2*) correlate positively with *LGR5*. **d**, Bowel sections from the human intestine were immunofluorescent stained for DEFA5 (red), LGR5 (green), and DAPI (blue). Scale bars, 50 μm in SI and 100μm in LI. Staining repeated on two to three participants (see also in Extended Data Fig. 9a-c). **e**, Expression of *LYPD8* and *WFDC2* in ST adult colon slides. H&E image of zoomed-in section repeated for clarity (see also in Extended Data Fig. 10a, b). **f**, Violin plots depicting selected AMPs genes (*DEFA5, LYPD8*, and *WFDC2*) that show locational and time-course differences in expression of fetal intestinal epithelium (top), and locational differences in the adult intestinal epithelium (bottom).

Further scRNA-seq profiles of fetal intestine^35^ showed that regional heterogeneity of AMPs appears at different time points before birth, such as 12-13 PCW for *DEFA5*, 15-16 PCW for *LYPD8* and 12-13 PCW for *WFDC2*, despite fetal AMP levels being lower than that of adults (Fig. 3f).

### The spatial distribution of AMPs along the crypt-villus axis suggested a unique niche of intestinal stem cells

Unexpectedly, the expression of *DEFA5/6, REG3A/G, ITLN2*, and *PRSS2* in small, especially proximal, intestine correlated positively with that of *LGR5* and decreased during cell maturation (Fig. 3a-c), indicating that stem cells (instead of Paneth cells which expressed the highest level of lysozyme-encoding gene *LYZ*) were the primary sources of certain AMPs^36,37^, such as defensins. Immunofluorescence demonstrated co-expression of *DEFA5*, *PRSS2*, and *LGR5* (Fig. 3d, Extended Data Fig. 9a-c). Laser capture microdissection and sequencing (LCM-seq) of mouse jejunal villus showed that AMPs enriched near the bottom of the villus (Extended Data Fig. 9d)^38^, which, together with our data, suggested that AMPs enriched in the lower part of the small intestinal crypt-villus axis. This distribution pattern of AMPs in SI might contribute to the maintenance of intestinal stem cell niche. In contrast, most colorectal AMPs were higher at the top of the crypt axis, as revealed by spatial transcriptomics (ST), except for *WFDC2* and *ITLN1* (Fig. 3e, Extended Data Fig. 10a, b).

Furthermore, ST analysis showed limited expression of AMPs in GALT (Fig. 3e, Extended Data Fig. 10a, b), where *CAMP* was mainly expressed by *CAMP*^+^ memory B cells while macrophages produced *HAMP*, *RNASE6*, and *LYZ* (Extended Data Fig. 9g-i). The articular structure of GALT might account for the unique AMP expression and help kill pathogens captured by microfold-like cells.

### Transcription factor regulatory network controls the regional expression of antimicrobial peptides

Accounting for the concurrent change of AMPs during epithelial differentiation (Fig. 3c), as well as the spatial co-expression patterns (Extended Data Fig. 10c), we hypothesized that there were standard upstream regulators for AMPs. To gain mechanistic insight, we generated transcription factor-target gene networks of SI- and LI-specific AMPs (Extended Data Fig. 11a, b), confirming the existence of common upstream TFs for AMPs. It is intriguing that SI AMPs, namely *DEFA5, DEFA6*, *PRSS2*, *REG3G*, *LEAP2*, and *REG3A*, were regulated by bile acid receptors *NR1H4* and *NR1H3* (Extended Data Fig. 11a), in line with the abundant expression of AMPs in *NR1H4*- and *NR1H3*-dependent intestinal stem cells, TA cells, and immature enterocytes (Fig. 2a). Consistently, the highest bile acid concentration in the ileum might lead to the elevated expression of AMPs in distal SI (Fig. 3a, b), highlighting the role of the microenvironment in innate immunity.

### A shift of AMPs in UC and Crohn’s disease (CD)

Abnormal expression of several AMPs has been noticed in patients with IBD^39–42^. To delineate a comprehensive AMP response during IBD, we analyzed the scRNA-seq data of noninflammatory and inflammatory biopsy of the colon from patients with UC, those of ileum from patients with CD, as well as those from healthy controls^12,43^. Colonic epithelium of patients with UC showed significant up-regulation of almost all AMPs, including SI-enriched AMPs, such as *DEFA5/6, REG3A*, and *ITLN2*, and the levels increased with the progression of the disease (healthy < non-inflammatory < inflammatory) (Extended Data Fig. 12a). In contrast, AMP alteration in the ileal epithelium of patients with CD showed an inconsistent trend (Extended Data Fig. 12a), suggesting distinct roles of AMPs in the pathogenesis of UC and CD. The increased AMPs during UC might be a compensatory response to the pathogenic microenvironment, whereas the suppression of AMPs even before endoscopic inflammation emerges during CD might exacerbate the condition.

Of note, a group of epithelial cells, which expressed goblet cell markers (*MUC2* and *ITLN1*), in the colonic epithelium of patients with UC showed up-regulated SI-specific AMPs (*DEFA5, DEFA6*, *REG3A*, and *ITLN2*) (Extended Data Fig. 12b, c). According to functional analysis of goblet cell subsets (Fig. 2h-k, Extended Data Fig. 7b-d), the AMP-producing cells were *TFF3*^+^ goblet cells (*TFF1*^−^*LYPD8*^−^) at the bottom of the crypt.

### Regional heterogeneity of adaptive immune surveillance

ScRNA-seq data revealed that plasma cells mainly populated the proximal SI and colon, whereas follicular B cells populated the ileum, consistent with the enrichment of Peyer’s patches (Extended Data Figs. 1e, 9e, f). The expression of IgA-related genes, such as *IGHA1* and *IGHA2*, was significantly higher in colonic plasma cells than in the small intestine. In contrast, IgM-related gene, *IGHM*, showed the opposite pattern (Extended Data Fig. 13a-c), indicating regional immune surveillance mediated by IgM/IgA class switch. Genes related to immunoglobulin synthesis, processing, and secretion also diverged between SI and LI (Extended Data Fig. 13d).

### Regional cores and heterogeneity of intestinal compartment/cell crosstalk

Cell-cell interaction^44^ and network representation analysis based on receptor-ligand pairs revealed that cell-cell crosstalk in different compartments represented the coexistence of compartmentalization and de-compartmentalization. More vigorous communication intensity display within the epithelial, fibroblast, and myeloid compartments (i.e., compartmentalization), whereas the T/ILCs, B, glia, and endothelial cells were de-compartmentalized (Fig. 4a-d). Significant regional heterogeneity was observed in the intensity of compartment/cell crosstalk (ileum > proximal SI > colon) (Fig. 4a-d), suggesting the ileum-predominant occurrence of CD, which in turn, is characterized by transmural inflammation, and penetrating and fibrotic stenosis, in contrast to the mucosal limitation in UC. The core of inter-compartment crosstalk in SI consisted of monocyte, macrophages, and stromal cells, whereas LI consisted of MNPs and epithelial cells (Fig. 4e).

**Fig. 4:**
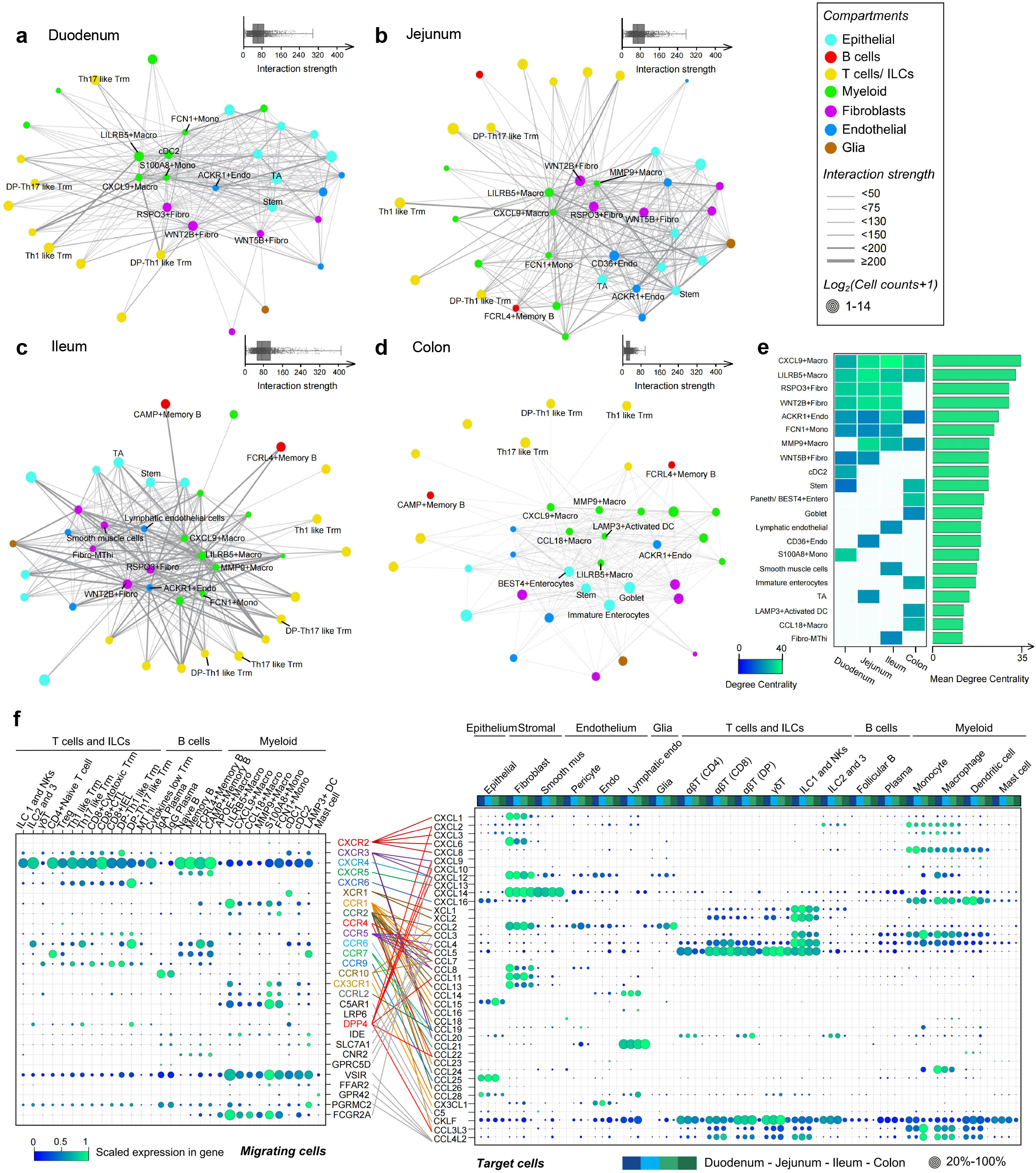
Regional heterogeneity of intestinal cell crosstalk. **a-d**, Cell-cell interaction networks based on receptor-ligand pairs in different intestinal regions including duodenum (a), jejunum (b), ileum (c), colon (d). Nodes, cell subsets annotated by lineage (color), and cell count (size). Edges connect pairs of cell subsets labeled by interaction strength (width). **e**, Central nodes in the interaction network. Shown are the top 10 central subsets of each intestinal region (left) and their mean degree centrality (connectivity) across different regions (right). **f**, Chemokine signaling predicts immune cell homing in different intestines. Dot plot showing expression of chemokine receptors (left) in immune cells and ligands (right) in intestinal cell compartments across different intestinal regions (colors). Only cell types and chemokines with detected expression are shown. Colored lines connect receptor sources (migrating cells) with target cell compartments expressing cognate ligand. Expression values are normalized and scaled. (see also in Extended Data Fig. 14).

### Molecular basis of intestinal immune cell homing

Chemokine-receptor pairs constitute the molecular basis for the competitive homing of immune cells. Results proved that the same lineage showed the region-specific ability of immune cell recruitment (Fig. 4f, Extended Data Fig. 14a, b), resulting in the spatial distribution of immune cells. For example, the increase of *CXCL1, 2*, and *3* in the epithelium, and their decrease in fibroblasts, toward the distal digestive tract (Extended Data Fig. 14a, b), suggested that neutrophils might be enriched near lamina propria fibroblasts in SI but recruited to the colonic epithelium.

Region-specific chemokines were identified, such as known *CCL25* in SI epithelium^45^ and newly identified *CCL3L3* and *CCL4L2* in SI T cells, NK-ILC1, and MNPs (Fig. 4f). New interactions were predicted, i.e., *GPR42-CCL4L2* mediated mast cell-T cell/NK-ILC1 interaction, *CX3CR1*-mediated homing of *LILRB5*^+^ macrophage, and *CXCL9*^+^ macrophage to *CX3CL1*-expressing fibroblasts and endothelia, and *CCR4*-mediated Treg homing to *CCL22*-expressing *LAMP3*^+^ DC (Fig. 4f). Furthermore, positive feedback of self-recruitment existed in all immune cell lineages, except naive T cells, follicular B cells, and mast cells (Fig. 4f).

### Mapping cellular and regional foci of intestinal diseases

Genome-wide association studies (GWAS) identified the risk alleles for intestinal diseases. We determined the expression of 85 risk genes of 9 diseases in all 59 cell subsets to identify the cellular origin of the diseases, such as the CD risk gene *IL23R* in ILC2-ILC3 and the food allergy (FA) risk genes *MMP12* and *STXBP6* in *MMP9*^+^ macrophage and mast cells, respectively (Extended Data Fig. 15a). Gene set enrichment analysis indicated enriched disease-risk genes in specific lineages. For example, the CD, UC, internal tuberculosis, and celiac disease risk genes were enriched in MNPs, and FA genes were enriched in mast cells, endothelial cells, and follicular B cells (Extended Data Fig. 15b). Multiple diseases correlated with immune cell disorders suggesting a crucial role of abnormal immune surveillance in the occurrence of intestinal diseases. We further investigated the cellular and regional susceptibilities to five diseases whose risk genes were enriched in the immune compartment, consistent with the epidemiological characteristics (Extended Data Fig. 15c, d). In accordance with the regional tendency, CD risk genes, *SNX20, CARD9, IL27*, and *PRDM1* enriched in ileum *CXCL9*^+^ macrophage, and *IL10* enriched in ileum *FCN1*^+^ monocyte. TB risk genes, *TAB3, MICB, TAP2*, and *C6orf47* enriched in ileum *S100A8*^+^ monocytes (Extended Data Fig. 15c).

## Discussion

We presented a comprehensive cell and spatial atlas across intestinal space, consisting of 59 cell subsets, of which eight were newly identified. The new subsets with unique gene expression, segment distribution, and spatial localization characteristics were associated with inflammatory progression and microbial defense. A transitional state of ILCs was identified, implicating a potential polarization and its effects in gut diseases^46,47^.

The atlas provided new insights into milieu-regulated epithelial differentiation and the regional molecular and functional characteristics. It indicated that anatomical location to be vital for the differentiation of regional epithelial subsets by GRN analysis. The sensitivity of different epithelial cell GRNs to the microenvironment appears to be variable.

Regional and spatial heterogeneity of AMP-mediated immune surveillance throughout development and adulthood was identified, suggesting a highly variable pattern of AMPs between SI and LI, which might be involved in shaping distinct microbial communities of SI and LI. A distinct role of AMPs in stem cell niche maintenance in different regions was identified. AMPs were concentrated at the bottom of the crypt in SI, but were mainly located at the top of the colonic crypts. This difference in spatial distribution of SI and LI AMPs might accord with the distinct structure of mucus layer, which needs further investigation.

Our data revealed that *LGR5*^+^ stem cells in SI, especially proximal, expressed the highest level of SI-specific AMPs, represented by *DEFA5/6*. This result is surprising since intestinal epithelial-derived microbial defense mechanisms used to be largely attributed to Paneth cells^36^. A mice study likely mirrored these *LGR5*^+^ cells in our data to be precursors that are committed to mature into differentiated secretory cells^48^. Common upstream TFs controlled the AMPs in SI and LI, illuminating the concurrent change of AMPs during epithelial differentiation, as well as the spatial co-expression patterns. Of note, the bile acid receptors, *NR1H3/4*, were involved in regulating the expression of SI-specific AMPs, emphasizing the involvement of milieu signals in AMP-mediated intestinal immune surveillance^49,50^ and consistent with gradually established regional AMPs pattern during fetal development, since around 17 PCW was the key window from which fetal bile secretion started^51^. Intriguingly, a goblet subset at the bottom of the crypt contributed to the up-regulation of defensins during UC, rather than previous-believed Paneth cell metaplasia^39,40^.

Furthermore, the correspondence of cell focus of risk genes to diseases’ location susceptibility, along with distinct cell-cell crosstalk and spatial heterogeneity of immune cells in different gut segments, provided new insights into regional milieu-determined immune surveillance in intestinal diseases. Overall, our data revealed the regional milieu of human intestine determined molecular cues of immune surveillance, dictating gut homeostasis and disease, which may yield dividends for human medicine.

## Acknowledgments

We thank Translational Medicine Core Facility of Shandong University for consultation and instrument availability that supported this work. This work was financially supported by a National Key Research and Development Program of China (2020YFA0804400); the National Natural Science Foundation of China (81873550, 82070552, 82071854, 82070540 and 81770538); a Youth Interdiscipline Innovative Research Group of Shandong University(2020QNQT009); the Taishan Scholars Program of Shandong Province; and a National Clinical Research Center for Digestive Diseases supporting technology project (2015BAI13B07).

## Author contributions

Y.L., X.Z. and S.L. initiated, designed and supervised the project; Y.Y. and L.L. carried out human tissue collection; M.Z. and B.F. performed human tissue processing and scRNA-seq experiments; M.Z. performed flow cytometry and immunofluorescence validation in the human and mice tissue and data interpretation; Y.W. analysed single-cell RNA sequencing and spatial transcriptomics data and generated figures; Y.W., M.Z., J.Z. and H.G. contributed to data visualization; Y.W., Y.Y., L.L., M.Z., J.Z., B.F., Z.L., R.J. and J.Q. contributed to interpretation of the results; Y.W., Y.Y., L.L., M.Z. and S.L. wrote the original draft; and Y.W., Y.Y., L.L., M.Z., X.Z., S.L., Y.L., M.L. and X.Y. conducted the review and editing of the manuscript.

## Extended data figures and tables

**Extended Data Fig. 1 | Strategy for scRNA-seq and meta-information of human intestinal samples and cells. a**, Overview of study design for intestinal regional atlas. **b**, The metainformation of the newly generated and previously published scRNA-seq data. **c**, Cell clustering and annotation pipeline. Cell expression profiles were computationally clustered by graph clustering algorithm and clusters were then separated into tissue compartments based on the expression of compartment-specific markers (*KRT8* (Epithelial cells), *CD3D* (T cells/ILCs), *CD79A* (B cells), *IGFBP7* (Stromal cells), *IGSF6* (MNPs) and *TPSAB1* (Mast cells)), as shown for UMAP plot of intestinal cell expression profiles. Cells from each tissue compartment were then iteratively re-clustered until differentially-expressed genes driving clustering were no longer biologically meaningful. Cell cluster annotation was based on the expression of canonical marker genes from the literature, ascertained tissue location, and inferred molecular function from differentially-expressed genes. **d**, Heap map of pairwise Pearson correlations of the average expression profile of each cluster. **e**, Spatial transcriptome projection of marker genes for newly identified macrophage and follicular B subsets highlighting the enrichment of subsets in GALT or lamina propria (*LILRB5* for M02, *CXCL9* for M03, *CCL18* for M04, *MMP9* for M05, *FCRL4* for B05, and *CAMP* for B06). The black box in the H&E image (left) indicates the location of GALT.

**Extended Data Fig. 2 | Diversity of epithelial and stromal cells in different intestinal regions and summary of major compartments, epithelial and stromal subsets in the human intestine. a**, Overview of major compartment characteristics from human intestine. The compartment-specific TFs and functional genes were obtained by differential expression analysis (DEA). EPI, epithelial cells; MNPs, monocytes, macrophages, and dendritic cells. **b**, Diversity of epithelial cells in different intestinal regions and their characteristics. Left: Shown are stacked frequencies of each subset divided by the respective total compartment frequency estimated by scRNA-seq in each intestinal region. Right: Overview of epithelial cell subset characteristics. **c**, Diversity of stromal cells in different intestinal regions and their characteristics. Left: Shown are stacked frequencies of each subset divided by the respective total compartment frequency estimated by scRNA-seq in each intestinal region. Right: Overview of stromal cell subset characteristics. Note that for Extended Data Fig. 2b, c, the subset-specific TFs were obtained by regulon specificity score (RSS) and functional genes were obtained by differential expression analysis. +++ indicates that the cell subset is detected and the proportion of it in the epithelial compartment of the respective region is more than 30%, ++ indicates that is more than 10% but less than 30%, + indicates that is more than 1% but less than 10%, -/+ indicates that is less than 1%, and - indicates no detection of this cell subset. The cell subsets marked in red are newly identified.

**Extended Data Fig. 3 | Diversity of immune cells in different intestinal regions and summary of immune subsets in the human intestine. a-c**, Diversity of T cells/ILCs (a), B cells (b), and myeloid cells (c) in different intestinal regions and their characteristics. Left: Shown are stacked frequencies of each subset divided by the respective total compartment frequency estimated by scRNA-seq in each intestinal region. Right: Overview of T cell/ILCs, B cell and Myeloid cell subset characteristics. For Extended Data Fig. 3a-c, the subset-specific TFs were obtained by regulon specificity score (RSS) and functional genes were obtained by differential expression analysis. +++ indicates that the cell subset is detected and the proportion of it in the epithelial compartment of the respective region is more than 10%, ++ indicates that is more than 1% but less than 10%, + indicates that is more than 0.3% but less than 1%, -/+ indicates that is less than 0.3%, and - indicates no detection of this cell subset. The cell subsets marked in red are newly identified.

**Extended Data Fig. 4 | ILCs transition state characteristics and FACS validation in human and mouse intestines. a**, UMAP embedding plot of T cells/ILCs compartment from SI showing 13 major subsets including NK-ILC1 and ILC2-ILC3. **b**, Trajectory and partition-based graph abstraction of NK-ILC1 and ILC2-ILC3 lineages showing a transition state of ILCs. **c**, UMAP embedding plot of ILCs compartment from SI depicting the clustering result of re-clustering of ILCs into 3 subsets (right). **d**, Heap map showing the scaled expression of marker genes of NK, ILC1, ILC2, and ILC3 for 3 re-clustering subsets (fig. S5C) highlighting a transition state of ILCs. **e**, FACS analysis of IL-22 and AREG expression after gating on CD45^+^Lin^-^ CD127^+^CD161^+^ ILCs and CD45^+^Lin^-^CD127^+^CD161^-^ LPLs in human SI (upper) and LI (lower) intestine. **f**, FACS analysis of IL-22 and AREG expression after gating on CD45^+^Lin^-^ CD127^+^CD90.2^+^ ILCs in mouse SI (upper) and LI (lower) intestine. **g**, The frequency of doublepositive ILCs was much higher in humans than mouse in both SI and LI. Data are shown as mean ± SEM (n =3 per group).

**Extended Data Fig. 5 | Transcription factor regulatory networks in scRNA-seq of healthy intestine. a**, UMAP embedding based on the TF module scores shows the intestinal cells into identical but consecutive cell lineages. **b**, UMAP overlay of selected TF module AUC scores in single cells across all compartments, demonstrating gene modules with compartment-specific regulation of intestinal epithelium (*CD2X*), T cells/ILCs (*STAT4*), B cells (*POU2F2*), MNPs (*SPIC*), mast cells (*GATA1*) and stromal cells (*TCF21*). **c**, Binary regulon activity heat map for the intestinal cell atlas. Binary activity for each cell is generated from the SCENIC AUC distribution and plotted as a heat map, with black blocks representing cells that are ‘on’. **d**, A dendrogram of regulons for 59 intestinal cell subsets. TFs at each branching point of the dendrogram are representative of subjacent groups of regulons. Branches are colored by lineages (from left to right, they are epithelium, endothelium, fibroblasts, glial, pericytes and smooth muscles, mast cells, monocytes-macrophages-dendritic cells, plasma cells, follicular B cells, and T/ILCs).

**Extended Data Fig. 6 | Overview of molecular and functional characteristics of regional and functional enterocyte subsets. a**, Functional characteristics of 6 functional enterocyte subsets. **b**, Dot plot showing the up-regulation of various mucus-related genes suggesting that enterocytes converged to goblet cells in mucus biosynthesis and secretion toward the end of the digestive tract, **c-e**, Function enrichment of 6 regional enterocyte subsets. Shown are normalized scores of KEGG pathways in organismal systems (c), metabolism (d), and human disease (e).

**Extended Data Fig. 7 | Overview of molecular and functional characteristics of regional and functional goblet subsets. a**, Functional characteristics of 4 functional goblet subsets. **b-d**, Trajectory and partition-based graph abstraction showing the differentiation of goblet functional subsets (b) and their regional distribution (c) and marker genes (d). **e-g**, Function enrichment of 7 regional goblet subsets. Shown are normalized scores of KEGG pathways in organismal systems (e), metabolism (f) and human disease (g).

**Extended Data Fig. 8 | Overview of molecular and functional characteristics of regional and functional Paneth and Paneth-like cells (*BEST4*+enterocytes) subsets. a-c**, Molecular (a) and functional (b, c) characteristics of 6 functional subsets. Note that there is only one differential gene up-regulated in subset 4, so there is no enriched pathway for subset 4. **d-f**, Function enrichment of 5 regional subsets. Shown are normalized scores of KEGG pathways in organismal systems (d), metabolism (e), and human disease (f).

**Extended Data Fig. 9 | Multi-omics verification of the regional heterogeneity of antimicrobial peptides expressed on the crypt-villus axis and different intestinal regions. a**, Bowel sections from human intestine are immunostained for REG3G. Scale bars, 100 μm. **b**, Bowel sections from human SI are immunofluorescent stained for DEFA5 (red), LGR5 (green) and DAPI (blue). *LGR5*^+^ stem cells are not the only source of *DEFA5*, a part of *DEFA5* positive epithelial cells are negative for LGR5. **c**, Bowel sections from intestine are immunofluorescent stained for PRSS2 (red), LGR5(green) and DAPI (blue). Scale bars, 50 μm in SI and 100 μm in LI. Staining repeated on two to three participants. **d**, LCM-seq of mouse jejunum villi showing the antimicrobial peptide-encoding genes *Lypd8, Leap2, Dmbt1, Ccl28* were enriched at the bottom of villi. **e**, B cell subset distributions indicating the newly identified B06-*CAMP*^+^ Memory B are enriched in the ileum. Shown are stacked frequencies of each B cell subset divided by the total B cell frequency estimated by scRNA-seq in each intestinal region. **f**, UMAP embedding plot of B cells highlighting expression and distribution of marker gene *CAMP* for B06 subset. **g-i**, ST slides of the adult colon showing the enrichment of *CAMP* (g), *LYZ* (h), and *WFDC2* (i) in the GALT.

**Extended Data Fig. 10 | The spatial transcriptome of colon depicting the enrichment location of AMPs on the crypt axis. a**, ST slides of the adult colon highlighting the enrichment of most LI-specific AMPs at the top zone of LI crypt. **b**, ST slides of the adult colon showing the enrichment of *ITLN1* and *WFDC2* at the bottom of LI crypt. **c**, ST slides of adult colon showing the rare co-expression (spatial co-localization) of SI-specific AMPs *REG3A*, *DEFA5*, and *DEFA6*.

**Extended Data Fig. 11 | Transcription factor-antimicrobial peptide regulatory network. a**, Venn network plot of SI-specific AMPs and their TFs highlighting the common upstream such as *NR1H4* and *HNF4G*. **b**, Venn network plot of LI-specific AMPs and their TFs highlighting the common upstream such as *CREB3L1* and *PPARG*. **c**, Interactive Venn diagram indicating 35 TFs overlapping from the upstream TFs of SI (47 TFs)- and LI (110 TFs)-specific AMPs.

**Extended Data Fig. 12 | The shift of antimicrobial peptides in ulcerative colitis and Crohn’s disease. a**, Box plots depicting the shift of AMPs during the progression of UC and CD. Shown are the mean expression of AMPs in epithelial cells from healthy (blue), non-inflamed (green), and inflamed (red) biopsies (t-test independent samples with Bonferroni correction, \**p* = 0.05; \*\**p* = 0.01; \*\*\**p* = 0.001; \*\*\*\**p* = 0.0001); boxplots: 25%, 50% and 75% quantiles). **b**, UMAP embedding plots of epithelial cells from UC biopsies highlighting expression and distribution of SI-specific AMPs *DEFA5, DEFA6, REG3A* and *ITLN2*. **c**, UMAP embedding plots of epithelial cells from UC biopsies showing the co-expression of marker genes of goblet cell and SI-specific AMPs, and low expression of markers (*TFF1* and *LYPD8*) in the top zone of crypt indicating a new goblet subset locating at the bottom of crypt expresses SI-specific AMPs during UC.

**Extended Data Fig. 13 | Clustering and functional heterogeneity of B cells in different intestine regions. a**, Heap map showing the normalized expression of marker genes for B cell subsets. **b**, UMAP embedding plot of B cells from different intestinal regions grouped in 6 clusters (4 for follicular B cells and 2 for plasma cells). **c**, UMAP embedding overlay showing the location distribution across all B cell subsets (left) and distribution of immunoglobulin-related genes *IGHM*, *IGHA1*, and *IGHA2*. **d**, Box plots showing the mean expression of genes and pathway signatures implicated in immunoglobulin synthesis and secretion in each B cell subset from different intestinal regions (t-test independent samples with Bonferroni correction, \**p* = 0.05; \*\**p* = 0.01; \*\*\**p* = 0.001; \*\*\*\**p* = 0.0001); boxplots: 25%, 50% and 75% quantiles).

**Extended Data Fig. 14 | Intestinal epithelial and stromal cell regional expression patterns of chemokines. a**, Dot plots showing the mean scaled expression of chemokines in epithelial cells from different intestinal regions (left), and regional expression patterns of epithelial subsets (right). **b**, Dot plots showing the mean scaled expression of chemokines in stromal cells from different intestinal regions (left), and regional expression patterns of stromal subsets (right).

**Extended Data Fig. 15 | Mapping cellular origins of intestinal disease by cell-selective expression of disease genes. a**, Dot plot of scaled expression of intestinal disease risk genes (columns) (numbered, associated disease shown above) enriched in specific intestinal cell subsets (rows). Red, newly identified cell subset association of gene. **b**, Violin plots depicting the enrichment of risk gene sets of 8 intestinal diseases in specific cell lineages. **c**, Dot plot of scaled expression of risk genes (rows) enriched in immune cell subsets (columns) and intestinal regions (colors), highlighting cellular and regional susceptibility to the five diseases (Crohn’s disease, ulcerative colitis, intestinal tuberculosis, B-cell malignancies, and food allergy). **d**, Matrix plot of scaled expression of risk genes (rows) for all cells (left) and selected immune cells (right) (used in Extended Data Fig. 15c) showing regional enrichment (columns).

**Supplementary Table 1**: an Excel file containing participants details.

**Supplementary Table 2**: an Excel file containing key transcription factors of 59 cell subsets, and TFs at each branching point representing subjacent groups of regulons.

**Supplementary Table 3**: an Excel file containing key transcription factors of 41 regional epithelial cell subsets, and TFs at each branching point representing subjacent groups of regulons.

## Materials and Methods

### Single-cell RNA-seq and spatial transcriptomics datasets collected in this study

We conducted the study in two phases: In the discovery phase, we integrated our newly generated scRNA-seq data of mucosal epithelium and lamina propria of duodenum, jejunum, and ileum, and previously published ileum, colon, and rectal epithelial data (GEO: GSE125970)^19^, and colon epithelial and lamina propria data (Single Cell Portal: SCP259)^10^ to generate a comprehensive human intestinal single-cell atlas. In the promotion phase, we integrated noninflammatory and inflammatory mucosal biopsies data of ileal Crohn’s disease (available for download at gutcellaltas.org)^43^ and colon ulcerative colitis (GEO: GSE116222)^12^, and fetal development data of ileum and colon ranged from 8 to 22 post-conceptual weeks (PCW) (GEO: GSE158702), as well as spatial transcriptome (ST) data of adult colon (GEO: GSE158328)^35^ and LCM-seq data of mouse jejunum (GEO: GSE109413)^38^ to identify regional immune surveillance shifts during diseases and human development, and the spatial regularity of intestinal crypt-villus axis. It is worth noting that in the discovery phase, we randomly sampled the colonic scRNA-seq data from Single Cell Portal (SCP259)^10^ to 1/3 of the original to balance the cell numbers of the proximal small intestine, distal small intestine and colon. In addition, before integrating data from different sources, the gene symbol of all open-source human single-cell sequencing data is modified to the gene symbol of GRCh38.p13 human reference genome.

### Human specimens for single-cell experiment

We generated scRNA-seq profiles from 14 intestinal samples collected from 10 healthy donors recruited in Qilu hospital of Shandong University at the time of routine gastroscopy or enteroscopy or colonoscopy (see also in Supplementary Table 1). Healthy volunteers were individuals without gastrointestinal tumors, polyps, or other organic diseases and who were overall healthy with no underlying diseases such as hypertension and diabetes. All samples were obtained with informed consent, and the study was approved by the medical science research ethics committee of Qilu hospital of Shandong University (KYLL-202008-127-1). All relevant ethical regulations of the medical science research ethics committee of Qilu hospital were followed.

### Single-cell collection and sorting

Intestinal mucosa was freshly sampled from the duodenum, jejunum, ileum of the volunteers, and intestinal biopsies were washed in PBS to remove mucus and blood cells. Intestine samples were then incubated with shaking in PBS containing 10 mM EDTA and 20 mM HEPES at 37°C for 20 min. After shaking, crypts and villus fraction in the medium were mechanically detached, strained, washed, and centrifuged; the pellet was then resuspended in warm TrypLE Express (GIBCO) and digested to single epithelial cells at 37°C. To obtain the lamina propria cell compartment, the rest pieces were digested by shaking in 2 mL of 5% (v/v) FBS in RPMI medium containing DNase I (Sigma) (150 μg/ml) and collagenase IV (Sigma) (1.5K U/ml) at 37°C for 20-30 min. The digested tissue was homogenized by vigorous shaking and filtered through 100 μm cell strainer. After centrifugation, the pellets were harvested and resuspended in a complete cell medium. Single-cell suspensions of epithelial and lamina propria cell components were pelleted, washed, strained, and resuspended in FACS buffer. 7-aminoactinomycin D (7-AAD) was added just before flow sorting. 7-AAD-negative living cells of epithelial and lamina propria compartment were sorted for further single-cell mRNA-sequencing separately. Data for all sorted cells were recorded for later experiments.

### Library preparation and single-cell RNA sequencing

Cells were concentrated to 700-1000 cells/μL and loaded on GemCode Single Cell Instrument (10x Genomics; Pleasanton, CA, USA) to generate single-cell gel bead-in-emulsions (GEMs). Next, GEMs were subjected to library construction using Chromium Single Cell 3’ Reagent Kits v2 (10x Genomics; Pleasanton, CA, USA) according to the manufacturer’s instructions, the steps of which included incubation at room temperature, complementary DNA amplification, fragmentation, end repair, A-tailing, adaptor ligation, and sample index polymerase chain reaction. To be compatible with BGISEQ-500 sequencing platform, libraries conversion was performed using the MGIEasy Universal Library Conversion Kit (App-A) (Lot: 1000004155, BGI). Then the converted library was subjected to subsequent DNA circularization and rolling- cycle amplification to generate DNA nanoballs. Purified DNA nanoballs were sequenced using the BGISEQ-500 sequencing platform, generating reads containing 16 base pairs of 10xTM barcodes, 10 base pairs of UMIs, and 100 base pairs of 3’ complementary DNA sequences.

### Alignment, quantification, and quality control of single-cell RNA sequencing data

Droplet-based sequencing data were aligned and quantified using the CellRanger software (version 3.0.2 for 3’ chemistry) using the GRCh38.p13 human reference genome. Scanpy (version 1.7.1)^52^ python package was used to load the cell-gene count matrix and perform quality control for newly generated dataset and collected datasets. For each sample, after removing the mitochondrial (gene symbols start with MT-) and ribosomal Protein (gene symbols start with RP) genes, cells with fewer than 2000 UMI counts and 250 detected genes were considered as empty droplets and removed from the datasets. After that, genes expressed in fewer than three cells were discarded.

### Doublet detection

To exclude doublets, we applied Scrublet software (version 0.2.3)^53^ to identify artifactual libraries from two or more cells in each scRNA-seq sample, including newly generated dataset and collected datasets. The doublet score for each single cell and the threshold based on the bimodal distribution was calculated with default parameters (sim_doublet_ratio=2.0; n_neighbors=None; expected_doublet_rate=0.1, stdev_doublet_rate=0.02). All remaining cells and cell clusters were further examined to detect potential false-negatives from scrublet analysis according to the following criteria: (1) Cells with more than 8000 detected genes, (2) Clusters that expressed marker genes from two distinct cell types, which are unlikely according to prior knowledge (i.e. *CD3D* for T cells and *EPCAM* for Epithelial cells). All cells or clusters flagged as doublets were removed from further downstream analysis.

### Graph clustering and partitioning cells into distinct compartments

Downstream analysis included normalization (*scanpy.pp.normalize_total* method, target_sum=1e4), log-transformation (*scanpy.pp.log1p* method, default parameters), cell cycle score (*scanpy.tl.score_genes_cell_cycle* method, cell cycle genes defined in Tirosh et al, 2016^54^, feature regress out (*scanpy.pp.regress_out* method, UMI counts, percentage of mitochondrial genes and cell cycle score were considered to be the source of unwanted variability and were regressed), feature scaling (*scanpy.pp.scale method*, max_value=10, zero_center=False), PCA analysis (*scanpy.tl.pca* method, svd_solver=‘arpack’), batch-balanced neighbourhood graph building (*scanpy.external.pp.bbknn* method, n_pcs=20)^55^, leiden graph-based clustering (scanpy.tl.leiden method, resolution=1.0)^56^, and UMAP visualization (*scanpy.tl.umap* method)^57^ performed using scanpy (version 1.7.1)^52^. Clusters were preliminarily partitioned into 6 compartments, using marker genes found in the literature in combination with differentially expressed genes (*scanpy.tl.rank_gene_groups* method, method=‘Wilcoxon test’). Specifically, epithelial compartment was annotated using a gene list (*EPCAM*, *KRT8*, *KRT18*, *KRT19*, *PIGR*), T and ILCs compartment (*CD2*, *CD3D*, *CD3E*, *CD3G*, *TRAC*, *IL7R*), B cell compartment *(JCHAIN, CD79A, IGHA1, IGHA2, MZB1, SSR4*), MNPs compartment *(HLA-DRA, CST3, HLA-DPB1, CD74, HLA-DPA1, AIF1*), Mast cell compartment *(TPSAB1, CPA3, TPSB2, CD9, HPGDS*, *KIT*), and Stromal cell compartment (*IGFBP7*, *IFITM3*, *TCF7L1*, *COL1A2*, *COL3A1*, *GSN*). See Extended Data Figs. 2, 3 for differentially expressed genes in all compartments.

### Re-clustering and differential gene expression to define cell subsets

Re-clustering and differential gene expression analysis *(scanpy.tl.rank_genes_groups* method, method=‘wilcoxon’) were performed on each cell compartment to accurately identify cell types or subsets and characterize the differential genes of each subset (Fig. 1c, Extended Data Figs. 2, 3).

For the epithelial compartment, the results of preliminary re-clustering indicated that some lineages from different intestinal regions, such as absorptive enterocytes from proximal SI, distal SI, and LI, were independent of each other in UMAP and cannot be grouped into a cluster. Therefore, we respectively performed clustering and annotation on epithelial compartment of proximal SI, distal SI, colon, and rectum, revealing nine cell types and 14 regional subsets in the intestine epithelium (Fig. 1a) and their proportion distribution and differential genes (Extended Data Fig. 2b). The cell types included *LGR5*^+^stem cell, transient-amplifying (TA) cell, goblet cell, enteroendocrine cell (EEC), tuft cell, immature enterocyte, and enterocyte that coexist across all segments, as well as small intestine-specific Paneth cell and colorectum-specific *BEST4*+Enterocyte (Paneth-like cell)^19^. Consistent with the results of Wang et al.^19^, tuft cells had a lack of detection in the rectal epithelium. Moreover, in line with previous intestinal scRNA-seq reports^12,19,35^, our healthy human intestinal epithelial data also did not detect rare microfold (M) cell, which was reported to expand in the colon during ulcerative colitis^10^.

For T/ILCs compartment, the results of preliminary re-clustering indicated that cells from the colon and small intestine were heterogeneous in UMAP. What’s more, the number of colon cells was much less than that of the small intestine. Therefore, we clustered and annotated the T/ILCs of SI and colon, respectively. The process of hierarchical clustering and annotation based on marker genes (Extended Data Fig. 3a) was as follows: First, we distinguished between αβ T and non-αβ T cells. The non-αβ T cells were further classified into γδ T cells > NK-ILC1 and ILC2-ILC3. The αβ T cells were preliminarily grouped into CD4^+^ Naive/Tcm, Treg, CD4^+^ Trm, CD8^+^ Trm, and CD4^+^CD8^+^ Trm based on mRNA scores of gene modules related to immunoregulation (*TNFRSF18, TNFRSF4, TNFRSF9, TNFRSF1B, AC133644.2, CTLA4, TIGIT, ICA1, RHBDD2, MAGEH1, IL2RA, TBC1D4, BATF, IKZF2, FOXP3*), naive/central memory (*SELL*, *LEF1, SOX4, SC5D, CCR7, TOB1, NOSIP)*, CD8/cytotoxic (*CD8A*, *CD8B, DHRS3, GZMA, GFOD1, IFITM3, PRF1*, *KLRD1*, *GZMB*, *CCL5*, *NKG7*, *FGR*, *CD160*, *FCER1G*, *XCL2*, *XCL1*, *GNLY*, *EOMES*, *CMC1*, *DTHD1*, *AOAH*, *CLIC3*, *CTSW*, *KLRF1*, *KLRC2*, *KLRC1*, *PTGDR*, *MCTP2*, *CCL3*, *CCL4, CCL3L3, CCL4L2, MATK, MAPK1, IL2RB)*, and resident memory T cell *(JUN, KLF6, FOSB, PTGER2, FOS, SYTL3, SPRY1, ANKRD28, GPR171, PDE4D, JAML, IL7R, GLIPR1, CD69*, *NFKBIA*, *PPP1R15A*, *NFKBIZ*, *TNFAIP3*, *PTGER4*, *ANXA1*, *ID2*, *ATF3*, *MGAT4A*, *AC092580.4*, *KLRB1*, *RORA*, *IL18R1*, *STAT4*, *IFNGR1*, *PFKFB3*, *GPR65*). Then, CD4^+^ Trm was further divided into CD4^+^ Type 1 cytokines Trm (Th1 like Trm) and CD4^+^ Type 3 cytokines Trm (Th17 like Trm). CD8^+^ Trm was further divided into CD8^+^ Cytotoxic Trm, CD8^+^ CTL Tem, and CD8^+^ IEL. The CD4^+^CD8^+^ Trm was further divided into DP-Type 1 cytokines Trm and DP-Type 3 cytokines-Trm. In addition, there was a cluster in the small intestine that showed very high mitochondrial gene, low ribosomal gene patterns, and low TCR gene expression patterns, and highly expressed mRNA splicing function-related genes, such as *FUS*, *HNRNPA2B1*, *HNRNPH1*, *HNRNPU*, *SRSF7*, and *RBM5*. Cells in this cluster contained CD4 and CD8 T cells, indicating cells aggregated beyond the CD4/CD8 lineage with their special gene expression patterns, and were annotated as MT_hi T cells. Furthermore, a cluster of CD4^+^ Trm in the colon expressed very low Th1 and Th17 cytokines, and was annotated as cytokines low Trm.

MNPs and B cells compartment did not show the same regional heterogeneity as epithelial or T/ILCs compartments. Therefore, we integrated data from different regions and re-clustered it together. The MNPs compartment was preliminarily divided into monocytes, macrophages, and dendritic cells. Further, based on the differential genes, the monocytes were divided into *S100A8*^+^ subset and *FCN1*^+^ subset, and the macrophages were divided into *APOE*^+^ subset, *LILRB5*^+^ subset, *CXCL9*^+^ subset, *CCL18*^+^ subset, and *MMP9*^+^ subset, and dendritic cells were divided into cDC1, cDC2, and *LAMP3^+^* Activated DC. The B cells compartment was preliminarily divided into follicular B cells and plasma cells. The follicular B cells were further divided into Naive B, Memory B, *FCRL4*^+^ Memory B, and *CAMP*^+^ Memory B. And the plasma cells were divided into IgA plasma and IgG plasma.

For the stromal cell compartments, the endothelial and glial cells did not show strong regional heterogeneity. According to the differential genes, the endothelial cells from different regions were re-clustered together and divided into *ACKR1*^+^ subset, *CD36*^+^ subset, *RGS5*^+^ subset, and Lymphatic endothelial cells. The glial cells clustered individually. The fibroblast compartment lacked prior knowledge of clear lineage distinctions like epithelial cells. Therefore, despite significant regional heterogeneity within the fibroblast compartment, we integrated data from different regions and re-clustered them together. As a result, the fibroblast compartment can be preliminarily divided into fibroblasts, pericytes, and smooth muscle cells, and the latter two had weaker regional heterogeneity. According to DEG and regional enrichment (Extended Data Fig. 2c), the fibroblasts were further divided into *RSPO3*^+^*OGN*^+^ subset (mainly enriched in the SI), *RSPO3*^+^*OGN*^-^subset (mainly enriched in the colon) and *WNT2B*^+^ subset-1 (SI), *WNT2B*^+^ subset-2 (colon), as well as *WNT5B*^+^ subset with weak regional heterogeneity, and Fibro-MThi subset with very high mitochondrial gene expression patterns.

### Estimation of cell proportions

Because EPI and LP samples were separately processed and sequenced, and the number of EPI and LP samples also varied in different intestinal regions, we cannot directly compare the enrichment of cell subsets in different intestinal segments by comparing cell numbers among regions. Therefore, we designed a normalized cell proportion metric (*proportion*\*) and calculated the enrichment of cell subsets of regions in each cell compartment, including epithelial, T/ILCs, B cell, MNPs and Mast, and Stromal cell compartment. For each compartment, the calculation process was as follows:

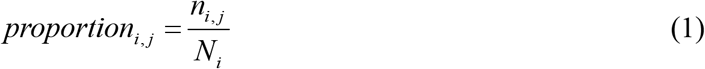

Where *i* and *j* indicated the intestine segment and cell subset, respectively. The proportion of cell subset *j* in the intestinal segment *i* is equal to the cell counts *n* of subset *j* in segment *i* divided by the total cell counts *N* in segment *i*.

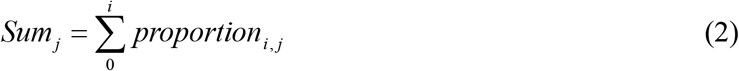

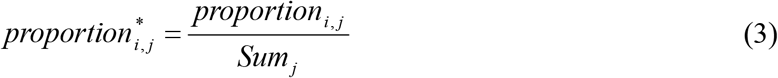

The normalized proportion (*proportion*\*) intuitively reflected the relative enrichment of cell types or subsets in each intestinal segment (Fig. 1f), and the proportion of each cell type in different segments added up to 1. If a cell type had the highest proportion in a particular intestinal segment, the proportion of this cell type in that intestinal segment exceeded other regions. The bar plots were done with matplotlib (version 3.3.4) and pandas (version 1.2.3) python package.

### Transcription factor module analysis

Python package pySCENIC workflow (version 0.11.0)^29,30^ with default settings was used to infer active TFs and their target genes in all cells and epithelial cells, respectively. In brief, the pipeline was implemented in three steps. First, a single-cell gene expression matrix was filtered to exclude all genes detected in fewer than ten total cells, and the remaining genes were used to compute a gene-gene correlation matrix for co-expression module detection using a regression per-target approach GRNBoost2 algorithm. Second, we pruned each module based on a regulatory motif near a transcription start site (TSS). Cis-regulatory footprints could be obtained with positional sequencing methods (for example, from ChIP-seq motif calling with an antibody against a TF). Binding motifs of TFs across multiple species were then used to build an RCisTarget database. Precisely, modules were retained if the TF-binding motif was enriched among its targets, while target genes without direct TF-binding motifs were removed. Third, we scored the impact of each regulon for each single-cell transcriptome using AUC score as a metric by the AUCell algorithm.

Each step of this pipeline used rank statistics, and the last classification step ran independently for each cell, avoiding a batch effect. The transcription factor motif scores for gene promoters and around transcription start sites for hg38 human reference genome were downloaded from the RcisTarget database^29^, and the TF gene list was downloaded from Humantfs database^58^. Based on the AUCell matrix of 543 regulons, dot-plot of top 3 activated regulons across 59 identified cell subsets and matrix plot of top 2 activated regulons across 41 regional epithelial cell subsets were done in the Scanpy package (*sc.pl.dotplot* method, standard_scale=‘var’) (*sc.pl.matrixplot* method, standard_scale=‘var’)^52^ (Extended Data Fig. 5d, Fig. 2A). GRN plots of the small intestine- and colorectum-specific antimicrobial peptides were done using the Evenn software^59^.

### Fate decision tree construction (regulon-based)

Dendrogram plots were constructed for all cells and epithelial cells using Scanpy (*sc.pl.dendrogram* method)^52^ on the AUCell matrix of 543 regulons to observe subtler changes, respectively (Figs. 1g, 2a, Extended Data Fig. 5d). We deciphered the diverging composite rules of a regulon-based dendrogram by testing each branching node for differential regulon importance. Therefore, we performed differential regulon expression analysis of every node with Wilcoxon test (*sc.tl.rank_gene_groups* method, method=‘Wilcoxon test’) to derive the action propagation program of the regulons. The dot plot of the fate decision tree was ordered by dendrograms (Extended Data Fig. 5d, Fig. 2A).

### Trajectory analysis

We applied the pseudo-time analysis to infer the differentiation trajectories of absorptive enterocytes and secretory cells across different intestinal regions (Fig. 2c, d), and then identified the influence of the regional microenvironment on the differentiation and functional shifts of epithelial cells. The analysis pipeline was implemented in two steps. First, clustering and annotation were performed on epithelial cells of the duodenum, jejunum, ileum, colon, and rectum, respectively (the methods were the same as <u>Graph clustering and partitioning cells into distinct compartments</u>). As a result, colonic enterocytes and goblet cells were divided into 2 and 3 functional subsets, respectively, and those in other regions had only one functional subset. In addition, considering the similarities between *BEST4*^+^ Enterocytes and Paneth cells in marker gene expression, such as *BEST4, HES4, SPIB, CA7*, and Lysozyme (*LYZ*), as well as growth factor receptor (*NOTCH2*), we classified these two cell types into one category (Panthe/*BEST4*^+^Enterocytes) and included them in the pseudo-time analysis of secretory cells. Second, we set *LGR5*^+^ stem cells as the initial point for differentiation and implemented pseudotime analysis to absorptive enterocytes and secretory cells (including functional subsets of goblet, tuft, EECs, Paneth/*BEST4*^+^ Enterocytes) with diffusion map (*sc.tl.diffmap* method, default parameters)^60^, Partition-based graph abstraction (PAGA) (*sc.tl.paga* method)^61^ and Force-directed graph drawing algorithm (*sc.tl.draw_graph* method, init_pos=‘paga’)^62^.

### Differentiation dynamics of antimicrobial peptides expression

We implemented the Wishbone python package (*scanpy.external.tl.wishbone* method, default parameters)^63^, an external module to the Scanpy package, to identify the dynamic trajectory of epithelial cell differentiation of proximal small intestinal and the expression of antimicrobial peptides along the trajectory (Fig. 3c). In brief, we processed the normalized gene-cell matrix in three steps. 1, the principal component analysis and batch correction based on BBKNN^55^ were performed. 2, we estimated the diffusion map of epithelium differentiation. 3, Wishbone^63^ and Phonograph^64^ python package were used to determine the differentiation branch and cluster the trends of antimicrobial peptides.

### Scoring gene set and identifying significant changes

We scored gene sets of all cells and clusters using the Scanpy python package (*sc.tl.score_genes* method, ctrl_size=len(genesets), gene_pool=None, n_bins=25, use_raw=None). The score was the average expression of a set of genes subtracted from the average expression of a reference set of genes. The reference set was randomly sampled from the gene_pool for each binned expression value. To prevent highly expressed genes from dominating a gene set score, we scaled each gene of the log_2_ (TP10K+1) expression matrix by its root mean squared expression across all cells. After obtaining the signatures score-cell matrix, differential signature analysis (*sc.tl.rank_gene_groups* method, method=‘Wilcoxon test’) was implemented to identify significant changes among different intestinal regions. All pathways used in gene set enrichment analysis of regional epithelial cells (Extended Data Figs. 6c-e, 7e-g, 10d-f) and B cells (Extended Data Fig. 13d) were obtained from KEGG^65^. In addition, we implemented gene set enrichment analysis to identify the distinct functions of epithelial cell subsets with metascape software^66^ (Fig. 2g, Extended Data Fig. 6a, 7a, and 8b, c).

### Cell-cell interaction and network representation analysis

To identify regional heterogeneity of cell-cell interaction, we applied the CellphoneDB python package^44^ (version 2.1.7) (*cellphonedb method statistical_analysis* method, threads=32, iterations=200) to the intestinal cell atlas, including duodenum, jejunum, ileum, and colon. Log transformed and normalized counts and cell subset annotations were used as an input. To narrow down the most relevant interactions, we looked for specific interactions classified by ligand/receptor expression in more than 10% of cells within a cluster and where log_2_ mean expression of ligand/receptor pair is greater than 0 *(cellphonedb plot heatmap_plot* method, default parameters). As a result, the file count_network output by the software recorded the count of interactions between cell subsets. To plot cell-cell interaction networks, we applied the Networkx (version 2.5) (https://github.com/networkx/networkx), Community (version 1.0.0b1) and, Pygraphviz (version 1.6) (https://github.com/pygraphviz/pygraphviz) python packages to a network defined using the count of interactions between cell subsets. For each network, the pipeline was implemented in three steps. First, we deleted the nodes whose degree is 0. Second, we removed the edges whose connection strength between nodes is less than the average strength of all edges. Third, we defined the size of nodes as log_2_ (counts+1) of cell subsets and characterized the network with the Kamada Kawai layout algorithm (*networkx.kamada_kawai_layout* method).

The chemokines-chemokines receptor interaction (Fig. 4F, fig. S17, and fig. S18) was obtained from IMEx Consortium^67^, IntAct^68^, InnateDB-All^69^, MINT^70^, and I2D^71^ database.

### Defining disease risk genes

We compiled lists of genes that have been implicated by human genetics studies as contributing to risk for the following diseases: Crohn’s disease (CD), Ulcerative colitis (UC), Intestinal tuberculosis (TB), Food allergy (FA), B-cell malignancies (BCM), Colorectal carcinoma (CRC), Irritable bowel syndrome (IBS), Celiac disease (CED) and Hirschsprung’s disease (HRSC). The risk genes curated from Genome-Wide Association Studies (GWAS) were obtained from the NHGRI-EBI Catalog (https://www.ebi.ac.uk/gwas/). And genome-wide association genes with log(p-value) < −20 or top 20 were included in our research.

### Flow cytometry and cell sorting of ILCs

Isolation of intestinal lamina propria cells was done as described. Lymphocytes isolated from human intestinal lamina propria were stained with antibodies against the following markers: CD45-PE/Cyanine7, CD127-BV421, CD117-PE/Cyanine5, CRTH2-APC/Cyanine7, CD161-Alexa Fluor 700, Amphiregulin, Rabbit IgG(H+L)-(R-PE), IL-22-APC. Lymphocytes isolated from mouse intestinal lamina propria were stained with antibodies against the following markers: CD45.2-Alexa Fluor 700, CD90.2-APC/Cyanine7, CD127-PE, IL-22-APC, Amphiregulin-Biotin, Streptavidin-eFluor 450. Lineage marker mix (Lin) for human contained FITC-CD3, CD19, CD123, CD14, CD303a, CD94, CD16. Lin for mouse contained FITC-FceR1, CD3, CD5, CD19, B220, Ly-6G, CD11b, CD11c, CD16/32 and TER-119. For cytokine staining, cells were stimulated with 50 ng/mL PMA, and 500 ng/mL ionomycin for 4 hr, and Brefeldin A (2 mg/mL) was added 2 hours before harvested. The live and dead cells were discriminated against by Zombie Aqua Fixable Viability Kit (BioLegend).

### Multi-color immunofluorescence

Human intestinal samples were fixed in 4% formaldehyde solution, dehydrated with ethanol, and embedded in paraffin. Four um-thick tissue sections on glass slides were deparaffinized through an ethanol gradient, and then tissue sections were incubated in retrieval solution for antigen retrieval at 95°C for 20 min. The sections were permeabilized and blocked for non-specific binding with 5% BSA and 0.1% Triton X-100 in PBS for 1 h at room temperature. Then, the sections were incubated overnight with the primary antibody at 4°C. The fluorescein-labeled secondary antibodies for immunofluorescence or secondary horseradish peroxidase-conjugated anti-rabbit antibody for immunohistochemistry were added for 1 h at 37°C. Slides were mounted with Slowfade Mountant+DAPI (Life Technologies, S36964) and sealed.

### Animal experiments

Animal experiments were carried out in compliance and approved by the Shandong university Specific Pathogen Free (SPF)-animal Center. All experimental animal procedures were approved by the Animal Care and Animal Experiments Committee of Shandong university (ECSBMSSDU2020-2-057). 6-8 weeks old male C57BL/6 mice were obtained from Nanjing GemPharmatech animal center and maintained in SPF facilities at Shandong University. Isolation of mouse intestinal lamina propria cells was done as previously described.

## Data availability

All raw and processed data will be deposited in the Gene Expression Omnibus (GEO).

## Code availability

The code generated during this study will be available at Github.

## References and Notes

1 Round, J. L. & Palm, N. W. Causal effects of the microbiota on immune-mediated diseases. Sci Immunol 3, doi:10.1126/sciimmunol.aao1603 (2018).

2 Atarashi, K. et al. Treg induction by a rationally selected mixture of Clostridia strains from the human microbiota. Nature 500, 232–236, doi:10.1038/nature12331 (2013).

3 Furusawa, Y. et al. Commensal microbe-derived butyrate induces the differentiation of colonic regulatory T cells. Nature 504, 446–450, doi:10.1038/nature12721 (2013).

4 Sassone-Corsi, M. & Raffatellu, M. No vacancy: how beneficial microbes cooperate with immunity to provide colonization resistance to pathogens. J Immunol 194, 4081–4087, doi:10.4049/jimmunol.1403169 (2015).

5 Buffie, C. G. & Pamer, E. G. Microbiota-mediated colonization resistance against intestinal pathogens. Nat Rev Immunol 13, 790–801, doi:10.1038/nri3535 (2013).

6 Britton, G. J. et al. Microbiotas from Humans with Inflammatory Bowel Disease Alter the Balance of Gut Th17 and RORgammat(+) Regulatory T Cells and Exacerbate Colitis in Mice. Immunity 50, 212–224 e214, doi:10.1016/j.immuni.2018.12.015 (2019).

7 Sartor, R. B. Microbial influences in inflammatory bowel diseases. Gastroenterology 134, 577–594, doi:10.1053/j.gastro.2007.11.059 (2008).

8 Schirmer, M., Garner, A., Vlamakis, H. & Xavier, R. J. Microbial genes and pathways in inflammatory bowel disease. Nat Rev Microbiol 17, 497–511, doi:10.1038/s41579-019-0213-6 (2019).

9 Kang, M. & Martin, A. Microbiome and colorectal cancer: Unraveling host-microbiota interactions in colitis-associated colorectal cancer development. Semin Immunol 32, 3–13, doi:10.1016/j.smim.2017.04.003 (2017).

10 Smillie, C. S. et al. Intra- and Inter-cellular Rewiring of the Human Colon during Ulcerative Colitis. Cell 178, 714–730 e722, doi:10.1016/j.cell.2019.06.029 (2019).

11 Martin, J. C. et al. Single-Cell Analysis of Crohn’s Disease Lesions Identifies a Pathogenic Cellular Module Associated with Resistance to Anti-TNF Therapy. Cell 178, 1493–1508 e1420, doi:10.1016/j.cell.2019.08.008 (2019).

12 Parikh, K. et al. Colonic epithelial cell diversity in health and inflammatory bowel disease. Nature 567, 49–55, doi:10.1038/s41586-019-0992-y (2019).

13 Zhang, L. et al. Single-Cell Analyses Inform Mechanisms of Myeloid-Targeted Therapies in Colon Cancer. Cell 181, 442–459 e429, doi:10.1016/j.cell.2020.03.048 (2020).

14 Pelka, K. et al. Spatially organized multicellular immune hubs in human colorectal cancer. Cell 184, 4734–4752 e4720, doi:10.1016/j.cell.2021.08.003 (2021).

15 Elmentaite, R. et al. Cells of the human intestinal tract mapped across space and time. Nature 597, 250–255, doi:10.1038/s41586-021-03852-1 (2021).

16 Maldonado-Contreras, A. L. & McCormick, B. A. Intestinal epithelial cells and their role in innate mucosal immunity. Cell and tissue research 343, 5–12, doi:10.1007/s00441-010-1082-5 (2011).

17 Graham, D. B. & Xavier, R. J. Pathway paradigms revealed from the genetics of inflammatory bowel disease. Nature 578, 527–539, doi:10.1038/s41586-020-2025-2 (2020).

18 Bunker, J. J. & Bendelac, A. IgA Responses to Microbiota. Immunity 49, 211–224, doi:10.1016/j.immuni.2018.08.011 (2018).

19 Wang, Y. et al. Single-cell transcriptome analysis reveals differential nutrient absorption functions in human intestine. J Exp Med 217, doi:10.1084/jem.20191130 (2020).

20 Morita, N. et al. GPR31-dependent dendrite protrusion of intestinal CX3CR1(+) cells by bacterial metabolites. Nature 566, 110–114, doi:10.1038/s41586-019-0884-1 (2019).

21 Picton, L. D. et al. A spinal organ of proprioception for integrated motor action feedback. Neuron 109, 1188–1201 e1187, doi:10.1016/j.neuron.2021.01.018 (2021).

22 Nighot, M. et al. Matrix Metalloproteinase MMP-12 promotes macrophage transmigration across intestinal epithelial tight junctions and increases severity of experimental colitis. Journal of Crohn’s & colitis, doi:10.1093/ecco-jcc/jjab064 (2021).

23 Yang, M. et al. Knocking out matrix metalloproteinase 12 causes the accumulation of M2 macrophages in intestinal tumor microenvironment of mice. Cancer Immunol Immunother 69, 1409–1421, doi:10.1007/s00262-020-02538-3 (2020).

24 Chen, Y. P. et al. Single-cell transcriptomics reveals regulators underlying immune cell diversity and immune subtypes associated with prognosis in nasopharyngeal carcinoma. Cell Res 30, 1024–1042, doi:10.1038/s41422-020-0374-x (2020).

25 Baumann, A., Demoulins, T., Python, S. & Summerfield, A. Porcine cathelicidins efficiently complex and deliver nucleic acids to plasmacytoid dendritic cells and can thereby mediate bacteria-induced IFN-alpha responses. J Immunol 193, 364–371, doi:10.4049/jimmunol.1303219 (2014).

26 Hurtado, P. & Peh, C. A. LL-37 promotes rapid sensing of CpG oligodeoxynucleotides by B lymphocytes and plasmacytoid dendritic cells. J Immunol 184, 1425–1435, doi:10.4049/jimmunol.0902305 (2010).

27 Jin, Y. et al. RGS5, a hypoxia-inducible apoptotic stimulator in endothelial cells. J Biol Chem 284, 23436–23443, doi:10.1074/jbc.M109.032664 (2009).

28 Kim, J. M., Kim, H. G. & Son, C. G. Tissue-Specific Profiling of Oxidative Stress-Associated Transcriptome in a Healthy Mouse Model. IntJMol Sci 19, doi:10.3390/ijms19103174 (2018).

29 Aibar, S. et al. SCENIC: single-cell regulatory network inference and clustering. Nat Methods 14, 1083–1086, doi:10.1038/nmeth.4463 (2017).

30 Van de Sande, B. et al. A scalable SCENIC workflow for single-cell gene regulatory network analysis. Nat Protoc 15, 2247–2276, doi:10.1038/s41596-020-0336-2 (2020).

31 Makishima, M. et al. Identification of a nuclear receptor for bile acids. Science 284, 1362–1365, doi:10.1126/science.284.5418.1362 (1999).

32 Modica, S., Gadaleta, R. M. & Moschetta, A. Deciphering the nuclear bile acid receptor FXR paradigm. Nucl Recept Signal 8, e005, doi:10.1621/nrs.08005 (2010).

33 Guzior, D. V. & Quinn, R. A. Review: microbial transformations of human bile acids. Microbiome 9, 140, doi:10.1186/s40168-021-01101-1 (2021).

34 Esterhazy, D. et al. Compartmentalized gut lymph node drainage dictates adaptive immune responses. Nature 569, 126–130, doi:10.1038/s41586-019-1125-3 (2019).

35 Fawkner-Corbett, D. et al. Spatiotemporal analysis of human intestinal development at single-cell resolution. Cell 184, 810–826 e823, doi:10.1016/j.cell.2020.12.016 (2021).

36 Bevins, C. L. & Salzman, N. H. Paneth cells, antimicrobial peptides and maintenance of intestinal homeostasis. Nat Rev Microbiol 9, 356–368, doi:10.1038/nrmicro2546 (2011).

37 Delorme-Axford, E. & Klionsky, D. J. Secretory autophagy holds the key to lysozyme secretion during bacterial infection of the intestine. Autophagy 14, 365–367, doi:10.1080/15548627.2017.1401425 (2018).

38 Moor, A. E. et al. Spatial Reconstruction of Single Enterocytes Uncovers Broad Zonation along the Intestinal Villus Axis. Cell 175, 1156–1167 e1115, doi:10.1016/j.cell.2018.08.063 (2018).

39 Cunliffe, R. N. et al. Human defensin 5 is stored in precursor form in normal Paneth cells and is expressed by some villous epithelial cells and by metaplastic Paneth cells in the colon in inflammatory bowel disease. Gut 48, 176–185, doi:10.1136/gut.48.2.176 (2001).

40 Ramasundara, M., Leach, S. T., Lemberg, D. A. & Day, A. S. Defensins and inflammation: the role of defensins in inflammatory bowel disease. J Gastroenterol Hepatol 24, 202–208, doi:10.1111/j.1440-1746.2008.05772.x (2009).

41 Wehkamp, J. et al. Reduced Paneth cell alpha-defensins in ileal Crohn’s disease. Proceedings of the National Academy of Sciences of the United States of America 102, 18129–18134, doi:10.1073/pnas.0505256102 (2005).

42 Courth, L. F. et al. Crohn’s disease-derived monocytes fail to induce Paneth cell defensins. Proceedings of the National Academy of Sciences of the United States of America 112, 14000–14005, doi:10.1073/pnas.1510084112 (2015).

43 Elmentaite, R. et al. Single-Cell Sequencing of Developing Human Gut Reveals Transcriptional Links to Childhood Crohn’s Disease. Dev Cell 55, 771–783 e775, doi:10.1016/j.devcel.2020.11.010 (2020).

44 Efremova, M., Vento-Tormo, M., Teichmann, S. A. & Vento-Tormo, R. CellPhoneDB: inferring cell-cell communication from combined expression of multi-subunit ligand-receptor complexes. Nat Protoc 15, 1484–1506, doi:10.1038/s41596-020-0292-x (2020).

45 Kunkel, E. J. & Butcher, E. C. Chemokines and the tissue-specific migration of lymphocytes. Immunity 16, 1–4, doi:10.1016/s1074-7613(01)00261-8 (2002).

46 Bielecki, P. et al. Skin-resident innate lymphoid cells converge on a pathogenic effector state. Nature 592, 128–132, doi:10.1038/s41586-021-03188-w (2021).

47 Goc, J. et al. Dysregulation of ILC3s unleashes progression and immunotherapy resistance in colon cancer. Cell, doi:10.1016/j.cell.2021.07.029 (2021).

48 Buczacki, S. J. et al. Intestinal label-retaining cells are secretory precursors expressing Lgr5. Nature 495, 65–69, doi:10.1038/nature11965 (2013).

49 Zhou, H. et al. Bile acid toxicity in Paneth cells contributes to gut dysbiosis induced by high-fat feeding. JCI Insight 5, doi:10.1172/jci.insight.138881 (2020).

50 Lu, R. et al. Paneth Cell Alertness to Pathogens Maintained by Vitamin D Receptors. Gastroenterology 160, 1269–1283, doi:10.1053/j.gastro.2020.11.015 (2021).

51 Colombo, C., Zuliani, G., Ronchi, M., Breidenstein, J. & Setchell, K. D. Biliary bile acid composition of the human fetus in early gestation. Pediatr Res 21, 197–200, doi:10.1203/00006450-198702000-00017 (1987).

52 Wolf, F. A., Angerer, P. & Theis, F. J. SCANPY: large-scale single-cell gene expression data analysis. Genome Biol 19, 15, doi:10.1186/s13059-017-1382-0 (2018).

53 Wolock, S. L., Lopez, R. & Klein, A. M. Scrublet: Computational Identification of Cell Doublets in SingleCell Transcriptomic Data. Cell Syst 8, 281–291 e289, doi:10.1016/j.cels.2018.11.005 (2019).

54 Tirosh, I. et al. Dissecting the multicellular ecosystem of metastatic melanoma by single-cell RNA-seq. Science 352, 189–196, doi:10.1126/science.aad0501 (2016).

55 Polanski, K. et al. BBKNN: fast batch alignment of single cell transcriptomes. Bioinformatics 36, 964–965, doi:10.1093/bioinformatics/btz625 (2020).

56 Traag, V. A., Waltman, L. & van Eck, N. J. From Louvain to Leiden: guaranteeing well-connected communities. Scientific reports 9, 5233, doi:10.1038/s41598-019-41695-z (2019).

57 Becht, E. et al. Dimensionality reduction for visualizing single-cell data using UMAP. Nat Biotechnol, doi:10.1038/nbt.4314 (2018).

58 Lambert, S. A. et al. The Human Transcription Factors. Cell 172, 650–665, doi:10.1016/j.cell.2018.01.029 (2018).

59 Chen, T., Zhang, H., Liu, Y., Liu, Y.-X. & Huang, L. EVenn: Easy to create repeatable and editable Venn diagrams and Venn networks online. Journal of Genetics and Genomics, doi:https://doi.org/10.1016/j.jgg.2021.07.007 (2021).

60 Haghverdi, L., Buttner, M., Wolf, F. A., Buettner, F. & Theis, F. J. Diffusion pseudotime robustly reconstructs lineage branching. Nat Methods 13, 845–848, doi:10.1038/nmeth.3971 (2016).

61 Wolf, F. A. et al. PAGA: graph abstraction reconciles clustering with trajectory inference through a topology preserving map of single cells. Genome Biol 20, 59, doi:10.1186/s13059-019-1663-x (2019).

62 Islam, S. et al. Characterization of the single-cell transcriptional landscape by highly multiplex RNA-seq. Genome Res 21, 1160–1167, doi:10.1101/gr.110882.110 (2011).

63 Setty, M. et al. Wishbone identifies bifurcating developmental trajectories from single-cell data. Nat Biotechnol 34, 637–645, doi:10.1038/nbt.3569 (2016).

64 Levine, J. H. et al. Data-Driven Phenotypic Dissection of AML Reveals Progenitor-like Cells that Correlate with Prognosis. Cell 162, 184–197, doi:10.1016/j.cell.2015.05.047 (2015).

65 Kanehisa, M., Furumichi, M., Tanabe, M., Sato, Y. & Morishima, K. KEGG: new perspectives on genomes, pathways, diseases and drugs. Nucleic Acids Res 45, D353–D361, doi:10.1093/nar/gkw1092 (2017).

66 Zhou, Y. et al. Metascape provides a biologist-oriented resource for the analysis of systems-level datasets. Nat Commun 10, 1523, doi:10.1038/s41467-019-09234-6 (2019).

67 Orchard, S. et al. Protein interaction data curation: the International Molecular Exchange (IMEx) consortium. Nat Methods 9, 345–350, doi:10.1038/nmeth.1931 (2012).

68 Orchard, S. et al. The MIntAct project--IntAct as a common curation platform for 11 molecular interaction databases. Nucleic Acids Res 42, D358–363, doi:10.1093/nar/gkt1115 (2014).

69 Breuer, K. et al. InnateDB: systems biology of innate immunity and beyond--recent updates and continuing curation. Nucleic Acids Res 41, D1228–1233, doi:10.1093/nar/gks1147 (2013).

70 Licata, L. et al. MINT, the molecular interaction database: 2012 update. Nucleic Acids Res 40, D857–861, doi:10.1093/nar/gkr930 (2012).

71 Brown, K. R. & Jurisica, I. Unequal evolutionary conservation of human protein interactions in interologous networks. Genome Biol 8, R95, doi:10.1186/gb-2007-8-5-r95 (2007).

