## Extended Figures for "Regional Cell Atlas of Human Intestine Shapes Distinct Immune Surveillance"

### Extended Data Fig. 1

**a** Study design, sample preparation, and sequencing

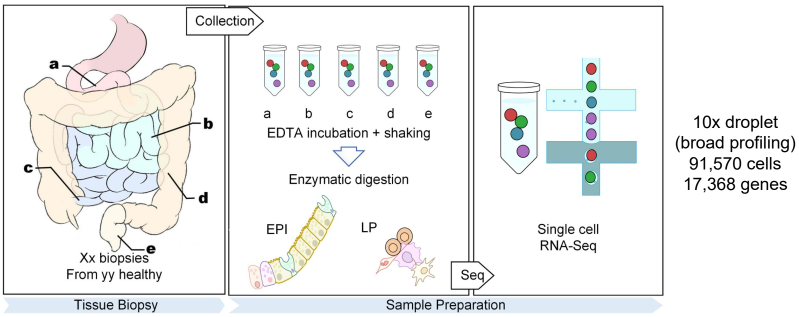


**b** Meta-information of the samples

| Organ | Tissue location | Number and source of samples | | | Number of cells after QC |
|---|---|---|---|---|---|
| | | Ours | SCP259 | GSE125970 | |
| Duodenum | Epithelium | 2 | 0 | 0 | 8276 |
| | Lamina propria | 3 | 0 | 0 | 11767 |
| Jejunum | Epithelium | 2 | 0 | 0 | 8149 |
| | Lamina propria | 3 | 0 | 0 | 13384 |
| Ileum | Epithelium | 2 | 0 | 2 | 7041 |
| | Lamina propria | 2 | 0 | 0 | 5132 |
| Colon | Epithelium | 0 | 24 | 2 | 17954 |
| | Lamina propria | 0 | 24 | 0 | 15992 |
| Rectum | Epithelium | 0 | 0 | 2 | 3875 |
| All | | 14 | 48 | 6 | 91570 |

**c** Cell clustering and annotation

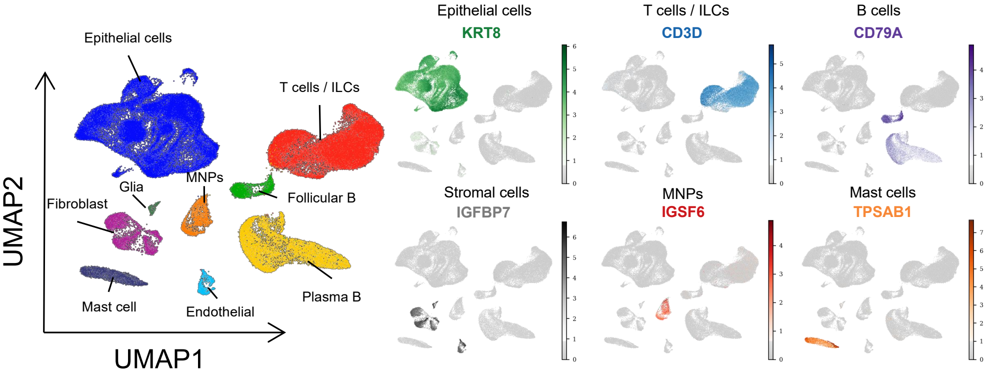


**d** Correlation between clusters

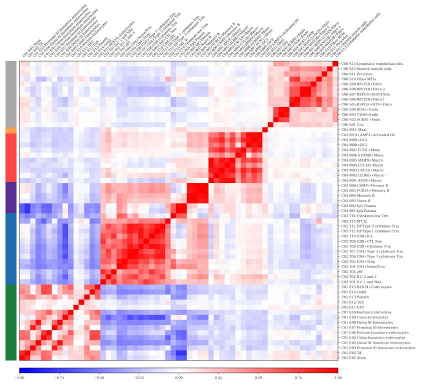

**e**

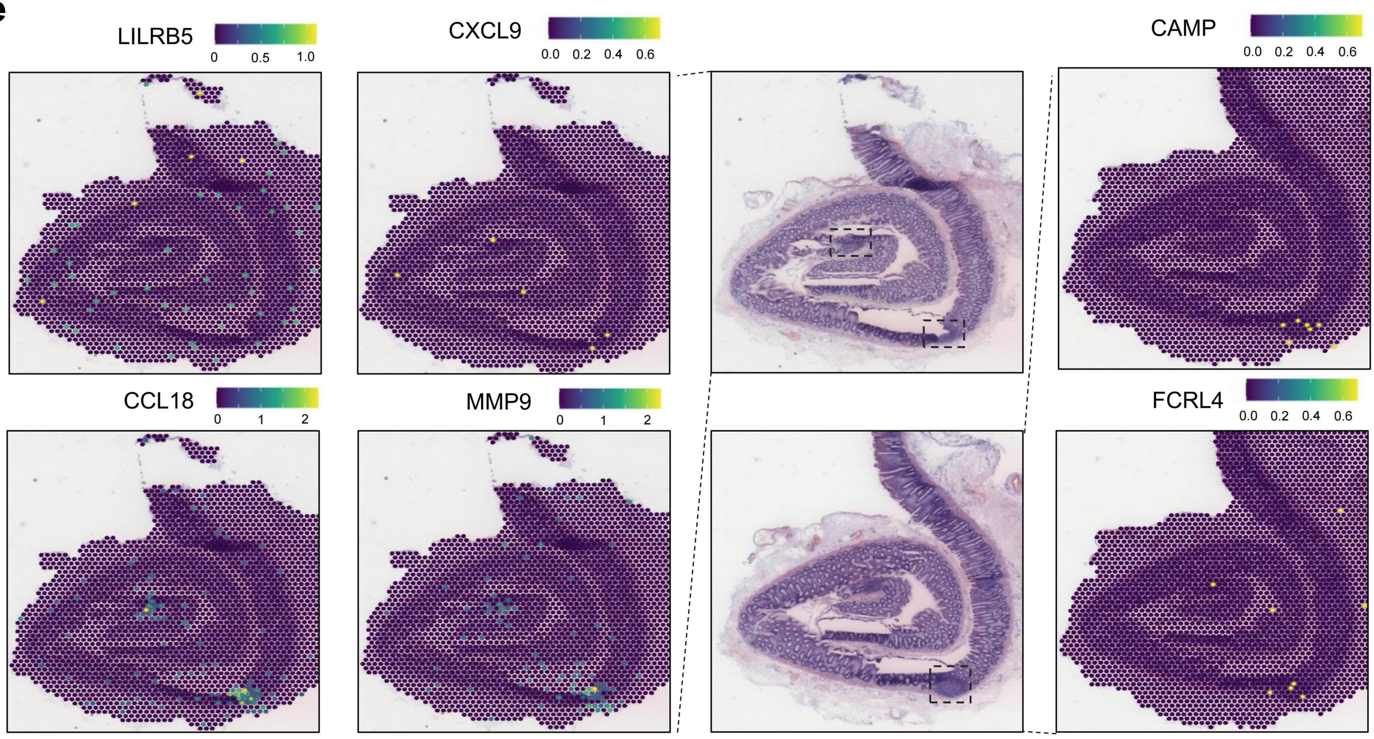

### Extended Data Fig. 2

**a**

| Human cell compartment | Representative genes | |
|---|---|---|
| | TFs (DEA) | other functional genes |
| **EPI** | FOXA3, CDX2, CDX1, GATA6, NR5A2 | PHGR1, KRT8, LGALS4, EPCAM, FABP1, KRT18, PIGR, TSPAN8, LGALS3, KRT19, CLDN3, CLDN7, ELF3, S100A6, C15orf48, CLDN4, SMIM22, S100A14, SPINK2 |
| **T** | TBX21, RORA, RUNX3, MYBL1, STAT4 | TRAC, IL32, CD3D, CCL5, CD2, TRBC2, CD3E, CD3G, EVL, IL7R, CD7, HCST, KLRB1, LCK, FYB1, CXCR4, CORO1A, |
| **B** | POU2F2, EBF1, E2F5, BACH2, RUNX2 | JCHAIN, CD79A, IGHA1, MZB1, DERL3, IGHA2, HERPUD1, SSR4, TNFRSF17, SEC11C, UBE2J1, PRDX4, GNG7, XBP1, EAF2, PLPP5, CD27, IGLC3, SSR3, TNFRSF13B |
| **MNPs** | ETV5, MAFB, IRF5, SPIC, ZNF467 | HLA-DRA, CST3, HLA-DPB1, CD74, HLA-DPA1, AIF1, LYZ, HLA-DQA1, C1QA, MS4A6A, HLA-DMA, C1QC, C1QB, FCGRT, DNASE1E3, LST1, SELENOP, FGL2, HLA-DMB, CTSB, GRN |
| **Mast** | GATA1, TAL1, NFE2L3, ZNF471, MSX2 | TPSAB1, CPA3, TPSB2, CD9, HPGDS, ANXA1, NFKBIA, MS4A2, CD63, LAPTM4A, SRGN, LMNA, LTC4S, FCER1G, VWA5A, CTSG, KIT, CLU |
| **Stromal cells** | TCF7L1, FOXF1, KLF9, TEAD2, RXRG, | IGFBP7, IFITM3, SPARC, A2M, CALD1, GSN, LGALS1, VIM, SPARCL1, CXCL14, COL1A2, COL3A1, COL6A2, TIMP1, C1S, S100A13, C1R, PLAT, MFAP4, RARRES2, COL1A1 |

**b**

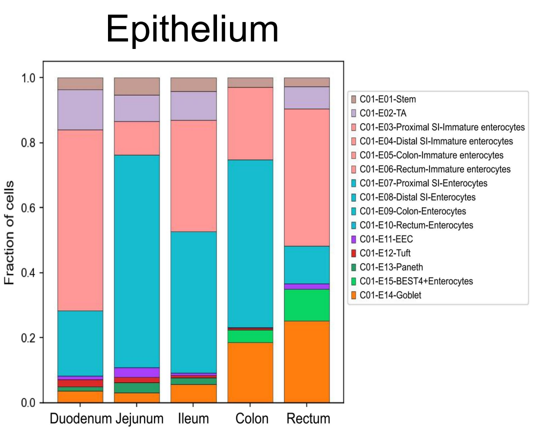


| Human epithelial cell cluster names (C01) | Shared markers | Representative genes | | Tissue enrichment | | | | |
|---|---|---|---|---|---|---|---|---|
| | | TFs (RSS top3) | other organ specific functional genes | D | J | I | C | R |
| **E01-Stem** | LGR5, RGMB, SMOC2, ASCL2 | DRGX, EVX2, POU4F1 | | + | + | + | + | + |
| **E02-TA** | MCM5, TOP2A, CCNA2, PCLAF | DRGX, ONECUT2, MNX1 | | ++ | + | + | - | + |
| **E03-Proximal SI-Immature enterosytes** | High expression of: OLFM4, DMBT1, CBR1, AKR7A3<br>Low expression of: organ specific functional genes in well differentiated enterosytes | GATA4, ONECUT2, PRDM16 | MTRNR2L12, SLC25A37, FOLH1, TMPRSS15, TM4SF4, TMEM238 | +++ | ++ | - | - | - |
| **E04-Distal SI-Immature enterosytes** | | NKX1-2, PITX2, GATA5 | FABP6, PRAP1, GUCA2A, LGALS2, DPEP1, DEFA5, CRIP1 | - | - | +++ | - | - |
| **E05-Colon-Immature enterosytes** | | DRGX, MECOM, TFCP2L1 | TMSB4X, C15orf48, ITM2C, FXYD3, FABP5 | - | - | - | ++ | |
| **E06-Rectum-Immature enterosytes** | | EVX2, HOXD13, FOXD2 | PRAC1, WFDC2, HOXB13, B3GNT7, MUC12, ST6GALNAC6 | - | - | - | - | +++ |
| **E07-Proximal SI-Enterosytes** | PHGR1, FABP1, GUCA2A, GUCA2B, SLC26A3 | GATA4, NR1I3, GATA5 | MTRNR2L12, TM4SF4, MTRNR2L8, FOLH1, CD36, MYO15B, C3orf85, ONECUT2, MIA2 | ++ | +++ | - | - | - |
| **E08-Distal SI-Enterosytes** | | NKX1-2, HES4, GATA5 | FABP6, GUCA2A, TFF3, FXYD3, ALDOA, ZG16, ITMXC, MT2A, DEFA5 | - | - | +++ | - | - |
| **E09-Colon-Enterosytes** | | TFCP2L1, ZFP30, ZFP64 | HMGCS2, MGST1, PEBP1, FABP5, UGT2B17 | - | - | - | +++ | - |
| **E10-Rectum-Enterosytes** | | BHLH41, TFCP2L1,TFAP2C | CD177, LGALS2, MUC12, SELENOP, CTSD, IVNS1ABP, CAMK2N1, SLC40A1, TFF1 | - | - | - | - | ++ |
| **E11-EECs** | PCSK1N, SCGN, CHGA, CACNA1A | NROB1, ISL1, PAX6 | | + | + | -/+ | -/+ | + |
| **E12-Tuft** | SH2D6, IRAG2, FYB1, BMX, AVIL | HMX3, ZFHX3, FOXP4 | | + | + | -/+ | -/+ | - |
| **E13-Paneth** | SPIB, CFTR, BEST4, ADGRG4, SULT1E1 | HES4, RBPJ, ONECUT2 | ALDOB, CFTR, MTRNR2L12, FAM3B, SI, REG1A, ADGRG4, FABP2, CPA2, OAT | + | + | + | - | - |
| **E15-BEST4+Enterocytes** | SPIB, BEST4, CA7, CA4, ITM2C | EVX2, FOXD2,HES4 | ALDOA, CA4, CA2, FABP5, CEACAM5, VSIG2, CKB | - | - | - | + | + |
| **E14-Goblet** | TFF3, FCGBP, KLK1, ITLN1, CLCA1 | ATOH1, EVX2, FOXD2 | | + | + | + | ++ | ++ |

**c**

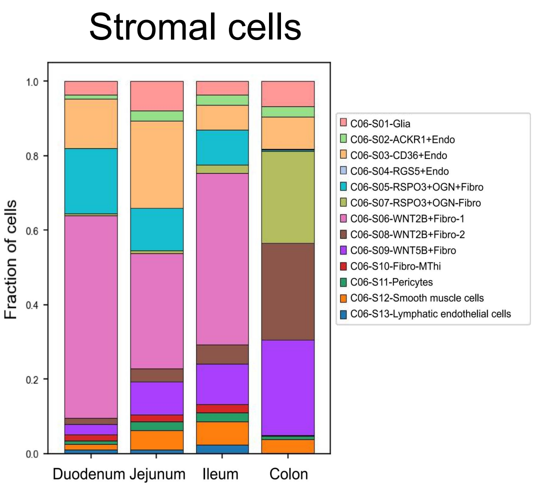


| Human stromal cell cluster names (C06) | Shared markers | | Representative genes | | Tissue enrichment | | | |
|---|---|---|---|---|---|---|---|---|
| | | | TFs (RSS top5) | other functional genes | D | J | I | C |
| **S01-Glia** | | | DLX5, DLX2, DLX1, ZNF740, SOX1 | CRYAB, ALDH1A1, S100B, GPM6B, CLU, PLP1, CD9, SPP1, NRXN1, SEMA3B, PRNP, LGI4, MPZ, PMEPA1, AP1S2, CDH19, MAL, FXYD1, | + | + | + | + |
| **S02-ACKR1+Endo** | PLVAP, CD320, FABP5, CD74, RAMP2, HLA-E, SRGN, IGFBP4, SPARCL1, RNASE1, JAM2, ESAM, VWF, IFI27, RAMP3, HRD4, ADGRL4, FLT1, CLEC14A | | LHX6, BCL6B, SOX17, FOXC1, NFIB | ACKR1, CPE, NPC2, CLU, MADCAM1, CLDN5, NNMT, DUSP23, ADGRG6, FBLN2, ADIRF, NR2F2, STXBP6, PRCP, KCTD12, ZNF385D, ICAM1, ACTN1, TIMP1, IL1R1, RAB3C, TPD52L1, IL33, PHACTR2, PERP, ALDH1A1, SELP, FAM155A, VEGFC | + | + | + | + |
| **S03-CD36+Endo** | | | BCL6B, NFIB, SOX17, EBF3, SMAD1 | GSN, CD36, VAMP5, PLVAP, SLC9A3R2, RGCC, CA4, PRSS23, MGP, TMEM88, FLT1, KDR, VWA1, TM4SF18, ACE, PLPP3, SLC14A1, PODXL, ADGRF5, MLEC, IGFBP3, INSR, GPRC5B | ++ | ++ | ++ | ++ |
| **S04-RGS5+Endo** | | | HEY1, SNAI1, ZNF214, ZNF155, MYCN | HIGD1B, RGS5, MYL9, DYNLRB1, SOD3, NDUFAL2, C12orf57, LGALS1, COX4I1, ATP5PO, NRN1, CCDC85B, FTH1, HLA-C, ARL6IP4, CIRBP, WARS1, PTP4A3, TMEM160, NDUFB2, LBHD1, TIMM8B, MIF, CHST12, GPX3 | - | -/+ | - | -/+ |
| **S05-RSPO3+OGN+** | CXCL14, MFAP4, CFD, C1S, DCN, C1R, RARRES2, LUM, COL3A1, FBLN1, APOE, CYGB, TCF21, ADAMDEC1, SERPINF1, COL6A2, COL1A2, MMP2, CTSC, COL1A1 | FBLN1, PTGDS, CCL11, PLAC9, CXCL12 | PRRX1, TWIST2, TP63, BHLHE22, PITX1 | JUND, IGKC, MARCKS, C11orf96, OGN, CD81, NFIA, TNC, COL6A3, ZBTB20, C7, CFH, ASPN, ZEB2, SELENOF, ANGPTL1, DDX17, IGLC2, CP | ++ | ++ | + | -/+ |
| **S07-RSPO3+OGN-** | | | TWIST2, PKNOX2, GLI1, PITX1, MSC | APOE, HLA-C, HLA-A, CCL8, ALDOA, WDR83OS, PDCD6, WFDC1, CXCL14, TXNDC12, FABP4 | -/+ | -/+ | + | +++ |
| **S06-WNT2B+Fibro-1** | | ADAMDEC1, CTSC, GGT5, PLPP3, ITIH5 | SOX15, PKNOX2, PITX1, PRRX1, MSC | C11orf96, JUND, ALDH1A3, CD81, MARCKS, CXCL6, HAS2, PDLIM1, CEBPD, SELENOF, BASP1, PLEKH2, ZEB2, ZBTB20, DACH1, CADM1, NR2F1, FILIP1L | +++ | +++ | +++ | - |
| **S08-WNT2B+Fibro-2** | | | PKNOX2, SOX15, MSC, PITX1, TCF21 | WDR83OS, ALDOA, S100A4, LSP1, U2AF1, APOE, SOD3, HLA-C, | + | + | + | ++ |
| **S09-WNT5B+Fibro** | | | ISL2, MSC, PITX1, GLIS2, FOXL1 | F3, CAV1, NSG1, PLAT, HSD17B2, FRZB, VSTM2A, ENHO, DMKN, EDNRB, POSTN, HLA-A, CD9, BMP4, PDGFD, SOX6, AGT, HLA-C, MMP11, MRPS6 | + | + | ++ | ++ |
| **S10-Fibro-MThi** | | | GLIS3, BCL6, TEAD1, NKX3-2, GLI1 | IKGC, IGLC2, IGHA1, JCHAIN, DDX17, DST, WSB1, MACF1, IGHM, ZEB2, VMP1, IGLC3, PNISR, HNRNPU, HNRNPH1, IGHA2 | + | + | + | -/+ |
| **S11-Pericytes** | ACTA2, TAGLN, MYL9, TPM1, TPM2, MYH11, ACTB, ACTG2, DSTN, MYLK, FLNA, RGS5, CKB, HHIP, SOSTDC1 | | HEY2, MEIS3, NR2F2, BCL6, EBF1 | IGFBP7, RGS5, NDUFA4L2, CD36, TINAGL1, GPX3, BGN, HIGD1B, NOTCH3, FABP5, PDGFRB, STEAP4, EBF1, COL18A1, EPS8, NR2F2, LHFPL6, ID3, FRZB, MCAM | -/+ | + | + | + |
| **S12-Smooth muscle cells** | | | MEIS3, MEF2D, GLI1, NR1D2, TEAD3 | CXCL14, ACTG2, TAGLN, HHIP, SOSTDC1, TPM2, PDLIM3, S100A6, CKB, MYL6, CD9, S100A13, NPNT, MFAP4, DCN, WFDC1, CNN1, S100A10, S100A4, HSD17B6, HSD17B6, LUM | + | + | + | + |
| **S13-Lymphatic endothelial cells** | | | ZNF746, EBF3, LHX6, BCL6B, SMAD1 | TFF3, LYVE1, CCL21, MMRN1, ECSCR, PROX1, FXYD6, GYPC, MARCKSL1, NRP2, PLIN5, ARL6IP1, RELN, MTUS1 | + | + | + | -/+ |

### Extended Data Fig. 3

**a**

T and ILCs

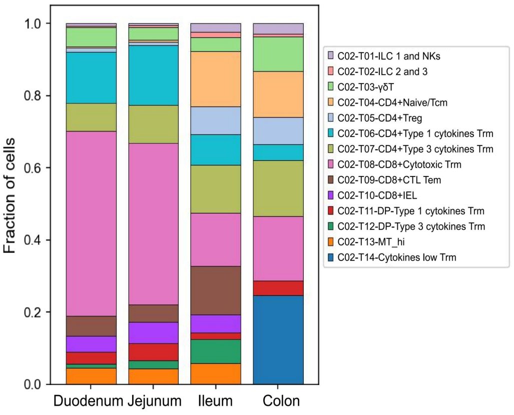


| Human T/ILCs cluster names (C02) | Shared markers | Representative genes | | Tissue enrichment | | | |
|---|---|---|---|---|---|---|---|
| | | TFs (RSS top5) | other functional genes | D | J | I | C |
| T01-ILC1 and NKs | High expression of: KLRB1, TYROBP, FCER1G<br>Low expression of: CD3D | GFI1, IKZF2, ZNF449, STAT4, TBX21 | TRDC, GNLY, GZMA/B/K, IFNG, PRF1, TNF, CCL3/4/5, XCL1, XCL2, IFGN, NKG7, KLRF1, KLRC1, ITGB2, CMC1 | + | + | ++ | ++ |
| T02-ILC2 and 3 | | ARNTL, TCF7, RORC, ZNF189, ZNF320, RORA | AREG, DLL1, IL2RA, KIT, IL7R, IL22, IL23R, CCR6, LTB, LST1, IL4I1, AQP3, TNFSF13B, MACROH2A1, ZFP36L1, TNFRSF25, AFF3 | -/+ | + | + | -/+ |
| T03-γδT | | GFI1, MYBL1, STAT4, IKZF2, TBX21 | TRDC, CCL5, ITGA1, TRAC, CD160, ITGAE, ENTPD1, ID3, ABI3, RAB3GAP1, CD247, CD7, CD3E, TRBC2, CDH17, KLHL23 | ++ | ++ | ++ | ++ |
| T04-CD4+Naive/Tcm | CD3D, CD4 | TCF7, LEF1, RBAK, ZFP64, TGIF2 | CCR7, KLF2, NOSIP, HLA-DRA, SELL, CD27, C1orf162, MAL, TRABD2A, TSHZ2, ACTN1, ARMH1, IL6ST | -/+ | + | ++ | ++ |
| T05-CD4+Treg | | FOXP3, BATF, RORA, LEF1, STAT4 | TIGIT, CD27, CTLA4, TBC1D4, ARID5B, BATF, TNFRSF4, TNFRSF4, CARD16, PTPRCAP, TNFRSF18, ICA1, DUSP4 | + | + | ++ | ++ |
| T06-CD4+Type 1 cytokines Trm | | TCF7, STAT4, MYBL1, RORA, GFI1 | CCL5, KLRB1, ITGA1, CCL4, TRGC2, CXCR3, CCR9, DTHD1, CCL2L2, METRNL, IFNG, MATK, IGFBP3 | ++ | +++ | + | + |
| T07-CD4+Type 3 cytokines Trm | | TCF7, STAT4, RORC, RORA, MYBL1 | IL4I1, CEBPD, CCR6, CCL20, TNFSF13B, PRR5, IFI44, IL17A, IL23R, IFI44L, DPP4, LTK, CTSH, CA2, SLC4A10, LST1, IL22 | ++ | ++ | ++ | ++ |
| T11-DP-Type 1 cytokines Trm | CD3D, CD4, CD8A | TCF7, MYBL1, STAT4, GFI1, RORA | **(Compared with T06)** HLA-C, IFITM1, MIF, CD160, CD7, CD8A, TMIGD2, H4C3, PLEKHF1, ABI3, CISH, CRIP1, XCL2, PSMB10 | ++ | ++ | + | ++ |
| T12-DP-Type 3 cytokines Trm | | TCF7, RORC, STAT4, MYBL1, RORA | **(Compared with T07)** MTRNR2L12, ITGA1, IKZF1, BCL11B, TXNIP, CXCR6, PHACTR2, IKZF3, MXD4, PLEKH1, GZMA, IFITM1, CD160 | + | ++ | ++ | - |
| T13-MT_hi | CD3D, CD4, CD8A | MYBL1, TCF7, ZNF66, KLF21, IKZF2 | N4BP2L2, DDX17, MBNL1, MTRNR2L12, NKTR, ARID1B, OGA, TTC14, OGT, LENG8, SMG1 | ++ | ++ | ++ | - |
| T14-Cytokines low Trm | CD3D, CD4 | ZFP64, ZNF2, ZNF684, ESR1, PRDM15 | ANXA1, PTPRCAP, ALDOA, NBEAL1, WDR83OS, LMNA | - | - | - | ++ |
| T08-CD8+Cytotoxic Trm | CD3D, CD8A | MYBL1, TCF7, GFI1, STAT4, IKZF2 | IL7R, KLRB1, S100A4, MGAT4A, FKBP11, CD9, JAML, SPINK2 | +++ | +++ | ++ | ++ |
| T09-CD8+CTL Tem | | STAT4, IKZF2, TBX21, MYBL1, EOMES | GZMK, GZMA, GZMH, KLRG1, NKG7, CD44, CCL3/4/5, HLA-DRA, CST7, CD81, APOBEC3G, CCL4L2, CMC1, SAMD3, HLA-DPB1 | ++ | ++ | ++ | - |
| T10-CD8+IEL | | GFI1, MYBL1, IKZF2, STAT4, TCF7 | KLRD1, CD7, CCL5, HOPX, CD160, CD63, ITGA1, KLRC2, TRGC1, METRNL, RAB3GAP1, IKZF2, ZEB2, FAM3C, ENTPD1, AKAP5, SLA2, AOAH, ADRB1 | ++ | ++ | ++ | - |

**b**

B cells

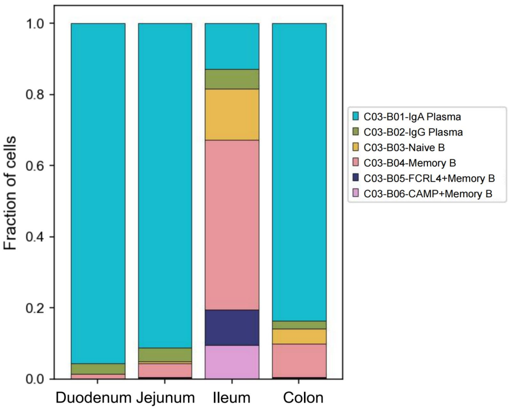


| Human B cell cluster names (C03) | Shared markers | Representative genes | | Tissue enrichment | | | |
|---|---|---|---|---|---|---|---|
| | | TFs (RSS top5) | other functional genes | D | J | I | C |
| B01-IgA Plasma | JCHAIN, SSR4, MZB1, FKBP1, IGHA1, IGHA2, TNFRSF17, XBP1, DERL3, SEC11C, PRDX4, SDF2L1, SSR3, NUCB2, DNAJB9, SELENOS, MANF, HSP90B1, TIMP1, CCR10, TNFRSF18 | ATF4, MEF2B, ZNF205, KLF11, CREB3L2 | IGHA1, IGHA2, TNFRSF17, JCHAIN, IGLL5, IGLC3 | +++ | +++ | ++ | +++ |
| B02-IgG Plasma | | ATF4, MEF2B, ZNF205, FOXN2, ZNF570 | IGHG1, IGHG3, IGHG4, IGHG2, IGHGP, VPREB3, RASSF6 | + | + | ++ | ++ |
| B03-Naive B | CD74, HLA-DRA, TMSB4X, PTMA, CXCR4, EEF1A1, CD52, ACTB, HLA-DPB1, BTG1, HLA-DPA1, MS4A1, CD37, LTB, CORO1A, VPREB3, LAPTM5, | POU2F2, BACH2, E2F5, ZNF320, ZNF296 | TCL1A, IGHD, FCER2, CD72, IL4R, YBX3, GPR18 | - | -/+ | ++ | ++ |
| B04-Memory B | | POUZF2, EBF1, BACH2, E2F5, RUNX2 | TNFRSF13B, GPR183, OCDC50, MARCKS, CD27, SAMSN1, AM2, TBC1D9, CLECL1, METTL8 | + | + | +++ | ++ |
| B05-FCRL4+Memory B | | POU2F2, RUNX2, MEF2B, EBF1, LYL1 | **(Compared with B04)** FCRL2/3/4/5, CD1C, ENTPD1, FAM107B, ACP5, CCR6, CTSH, ACTG1, IFNGR1, GSN, PTPN1, PLEKHO1, PXX, ITGAX, MYH9 | -/+ | -/+ | ++ | -/+ |
| B06-CAMP+Memory B | | POU2F2, EBF1, RUNX2, E2F5, BACH2 | **(Compared with B04)** CAMP, CD1C, FCRL2, GRN, PDCD4, GRB2, FCRL3, SELENOF, TSPAN33, CR2, GNAS, PPP1R14B, ACP5, SESN3, ACTR3, PPPAR2, CUL3, SET, CDV3, FCRLA | - | -/+ | ++ | -/+ |

**c**

MNPs and Mast cells

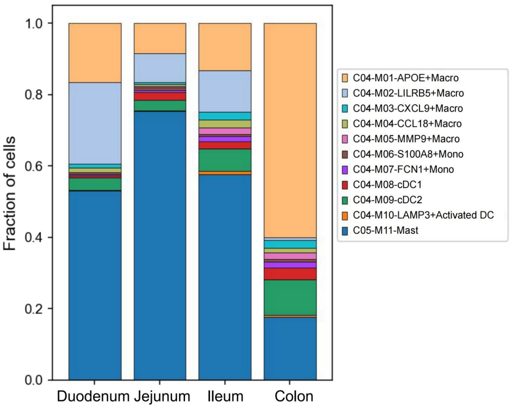


| Human myeloid cell cluster names (C04/05) | Shared markers | Representative genes | | Tissue enrichment | | | |
|---|---|---|---|---|---|---|---|
| | | TFs (RSS top5) | other functional genes | D | J | I | C |
| M01-APOE+Macro | SELENOP, C1QB, C1QC, C1QA, FTL, CTSD, SLC40A1, CTSB, CTSC, STAB1, LGMN, MS4A7, DAB2, PLD3, ITM2B, TMEM176B, IGD1, MAF, APOE, LIPA, CD14, A2M, CD63, FCGRT, CTSL, SLCO2B1, MS4A4A, CD209 | SPIC, ETV5, IRF5, ZNF467, MAFB | HLA-DRB5, APOE, RNASE1, APOE, WDR83O, LGALS3, APOC1, S100A4, FABP3, S100A9, S100A4, OTOA, | ++ | + | ++ | ++ |
| M02-LILRB5+Macro | | SPIC, ZNF732, ETV5, PLAGL1, IRF5 | **(Compared with M01)** CCL3L3, MARCHF1, N4BP2L2, AKAP9, GADD45B, DUSP6, KLF6, NAMPT, ZFP36L1, AHR, REL, JUND, CCNL1, FOS, ZEB2, ATP2B1, EIF4A1, UTRN, PNISR, ITPR2, CCDC88A, PTPRC, WSB1, VMP1, CCL4L2, OTUD1, MAFB, DDX17, CEBPD | ++ | + | ++ | -/+ |
| M03-CXCL9+Macro | | SPIC, IRF5, ETV5, MAFB, TFEB | **(Compared with M01)** CXCL9, CXCL10, SOD2, WARS1, TNFAIP2, STAT1, GBP1, KLF6, SRGN, PIM3, SLAMF7, GPR183, HNRNPC, IRF1, BCL2A1, GBP2, PPA1, ICAM1, GBP4, TANK, TNFSF10, BIRC3, NFKBIZ, DSE, SGK1, REL, PLEK, PTPRE, LAP3 | -/+ | -/+ | -/+ | -/+ |
| M04-CCL18+Macro | | SPIC, ETV5, MZF1, IRF5, TCF12 | **(Compared with M01)** CCL18, CTSC, SELENOP, SLC40A1, F13A1, FOLR2, LGMN, | -/+ | -/+ | -/+ | -/+ |
| M05-MMP9+Macro | | SPIC, IRF5, ETV5, TFEB, MAFB | **(Compared with M01)** MMP9, MMP12, CTSH, DNASE1L3, IL4I1, LGALS2, PLA2G7, TMEM176B, ITGB2, ENPP2, CD4, CTSA, CAPG, SELENOS,CFP | -/+ | -/+ | -/+ | -/+ |
| M06-S100A8+Mono | FCN1, S100A4, S100A6, TSPO, SOD2, S100A9, SERPINA1, IL1B, S100A8, C5AR1, PRKCB, CSTA, CD44 | NFE2, SPIC, GRHL1, IRF5, OTX1 | S100A8, S100A9, S100A11, S100A6, S100A4, S100A12, SLC11A1, CTSS, BRI3, STXBP2, MNDA, VCAN, LRRK2, MT2A, EVI2A, TALDO1, FGR, MBD2, RIPOR2, DUSP6 | -/+ | -/+ | -/+ | -/+ |
| M07-FCN1+Mono | | SPIC, NFE2, MZF1, SNAI1, IRF5 | HLA-DPB1, HLA-DPB1, HLA-DMB, HLA-DQA1, HLA-DMA, HLA-DQB1, HLA-DOA, HLA-DRB5, CD74, CLEC10A, EEF1B2, HSPE1, HMGN1, ZNF331, C1QA, GPR183, HERPUD1, P2RY6, RSU1, SLAMF8, C1QC, HES1, PLAAT4, MS4A6A | -/+ | -/+ | -/+ | -/+ |
| M08-cDC1 | LGALS2, LSP1, PPA1, BASP1, CD1E, FCER1A, CD1C, LTB, CST7, IDO1, PKIB, SLC38A1, RGCC | ZNF732, SPIC, TFEB, IRF5, HOXD9 | CLEC9A, RGCC, SNX3, C1orf54, IRF8, CADM1, ID2, RUBCNL, CPNE3, CCND1, GYPC, FUCA1, TMEM14A, AM2, GCSAM, CLNK, SUSD3, WDFY4, CXR1, VAC14, ENPP1 | -/+ | -/+ | -/+ | + |
| M09-cDC2 | | SPIC, IRF5, TEFB, ZNF732, SNAI1 | FCER1A, LST1, CLEC10A, JAML, FCER1G, TYROBP, IL1B, MS4A6A, IGSF6, CD1E, CD1C, RNF130, S100A4, PLD4, CFP, CASP1, S100A6, HCST, FCGR2B, PLAC8, CD300A, IFI30, DOK2, IFITM3 | + | + | + | ++ |
| M10-LAMP3+Activated DC | | FOXD4, LBX2, ZNF264, SPIC, ZNF732 | LAMP3, HLA-F, MARCKSL1, FSCN1, BIRC3, TMEM176A, TRAF1, GABARAPL2, MARCKS, GSN, CD44, CST7, KDM2B, TMEM176B, CCR7, GPR157 | -/+ | -/+ | -/+ | -/+ |
| M11-Mast | | NFE2, GATA1, NFE2L3, FOXI2, POU4F1 | **(Compared with MNPs)** TPSAB1, CPA3, CD9, TPSB2, HPGDS, MS4A2, CD69, GATA2, CTSG, VWA5A, LTC4S, LMNA, CLIC1, PPP1R15A, JUN, KIT, CLU, ALOX5AP | ++ | ++ | ++ | ++ |

### Extended Data Fig. 4

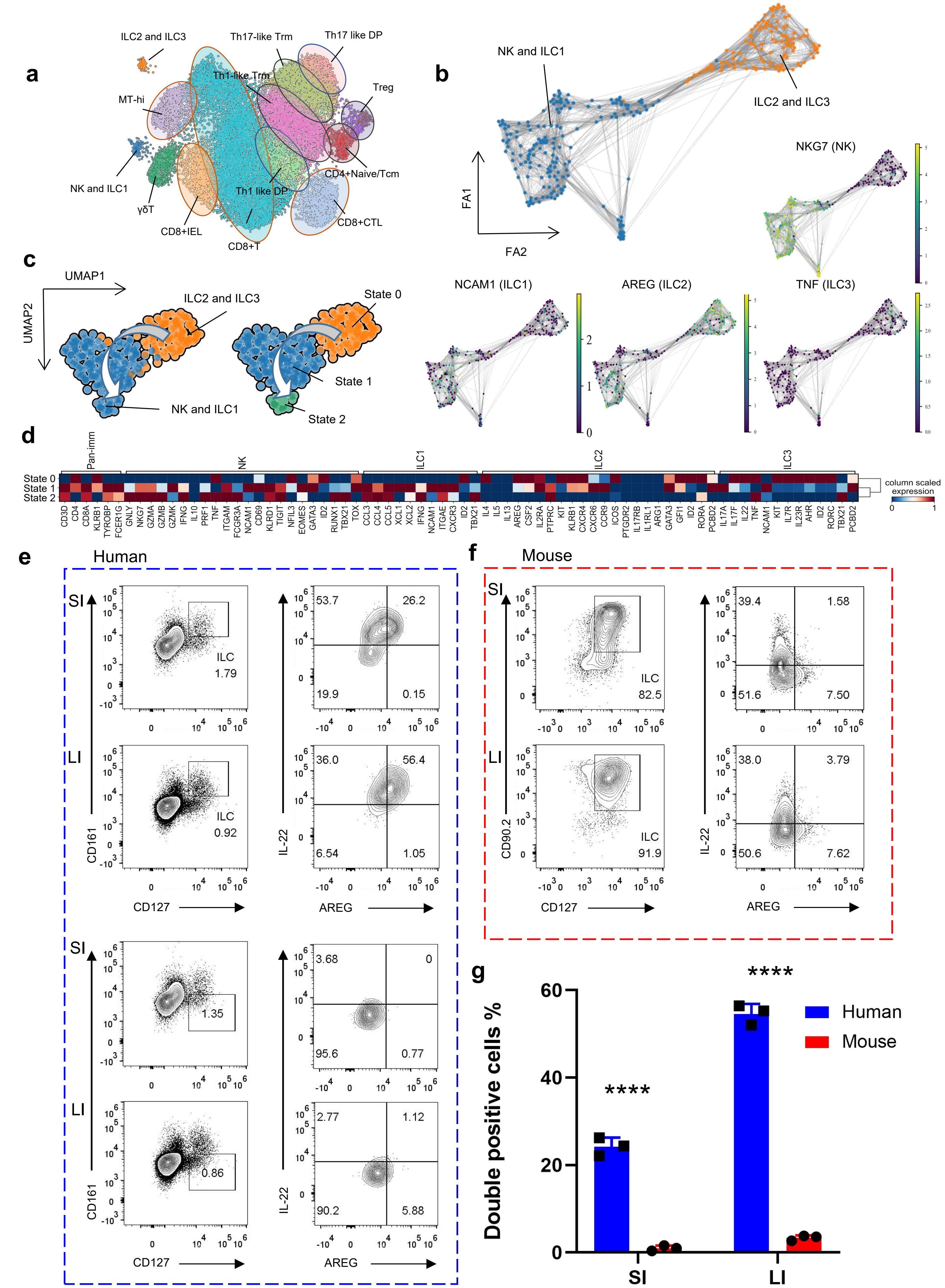

### Extended Data Fig. 5

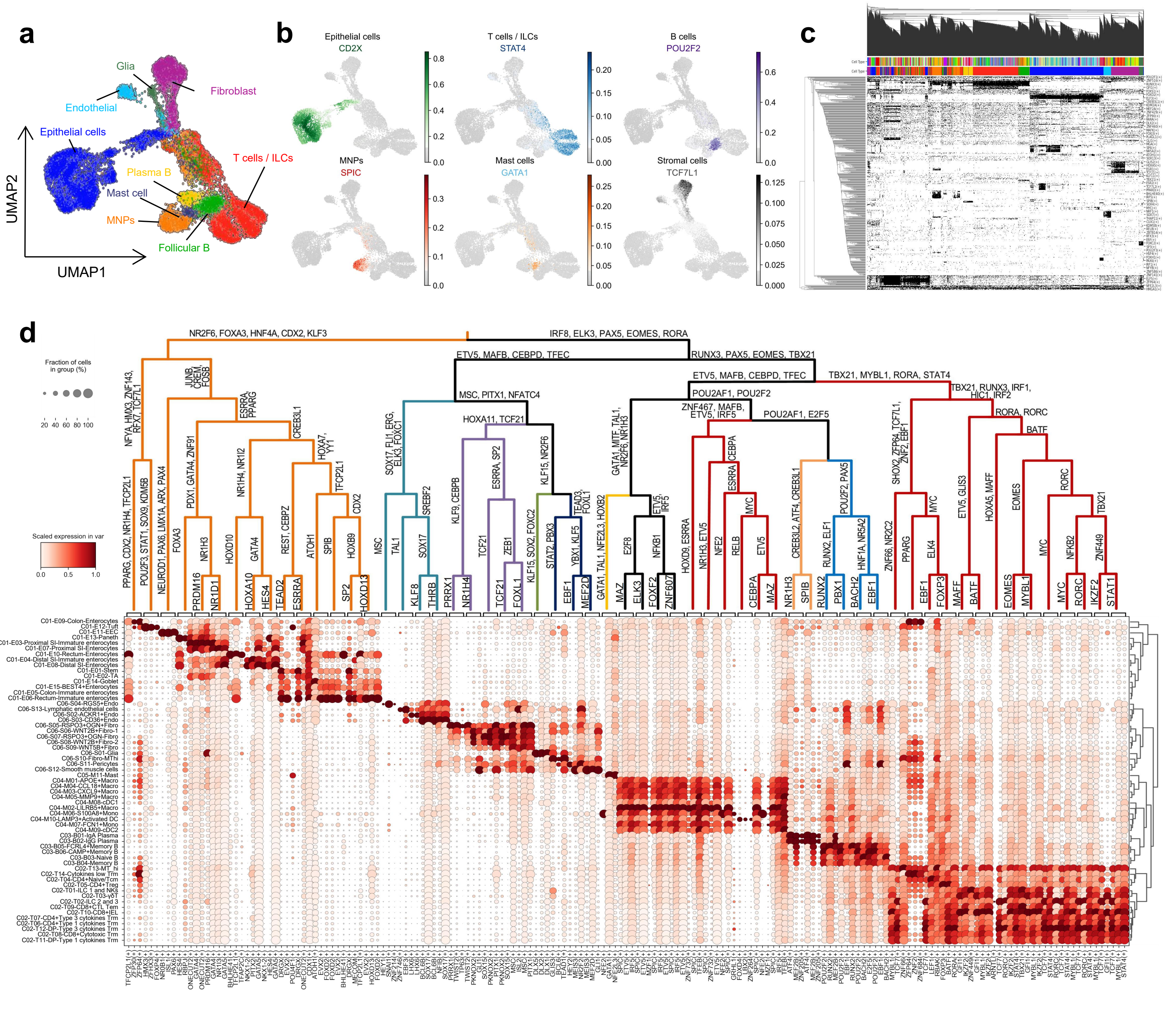

### Extended Data Fig. 6

Enterocyte

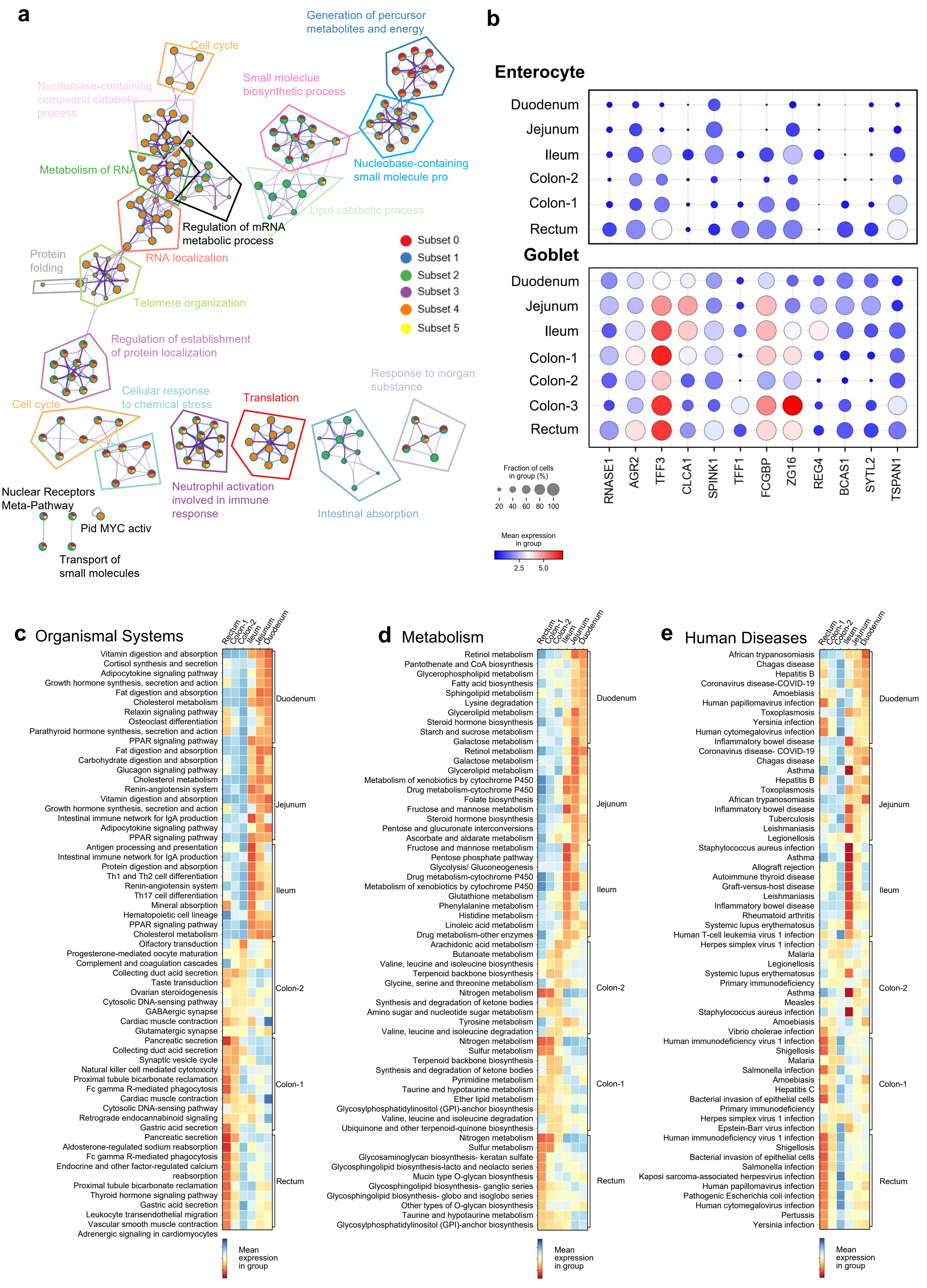

### Extended Data Fig. 7

Goblet

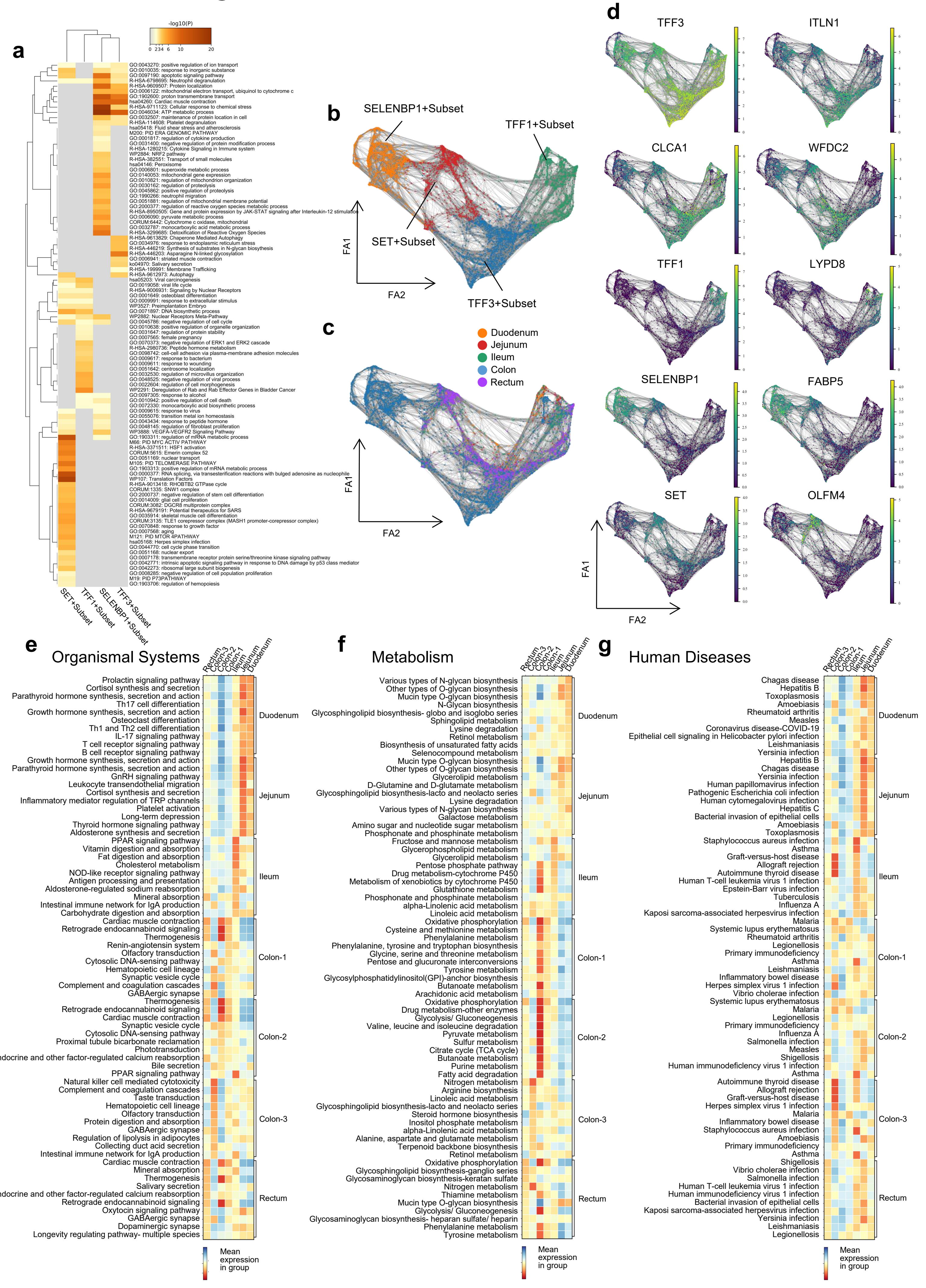

### Extended Data Fig. 8

Paneth and PLC (BEST4+Enterocytes)

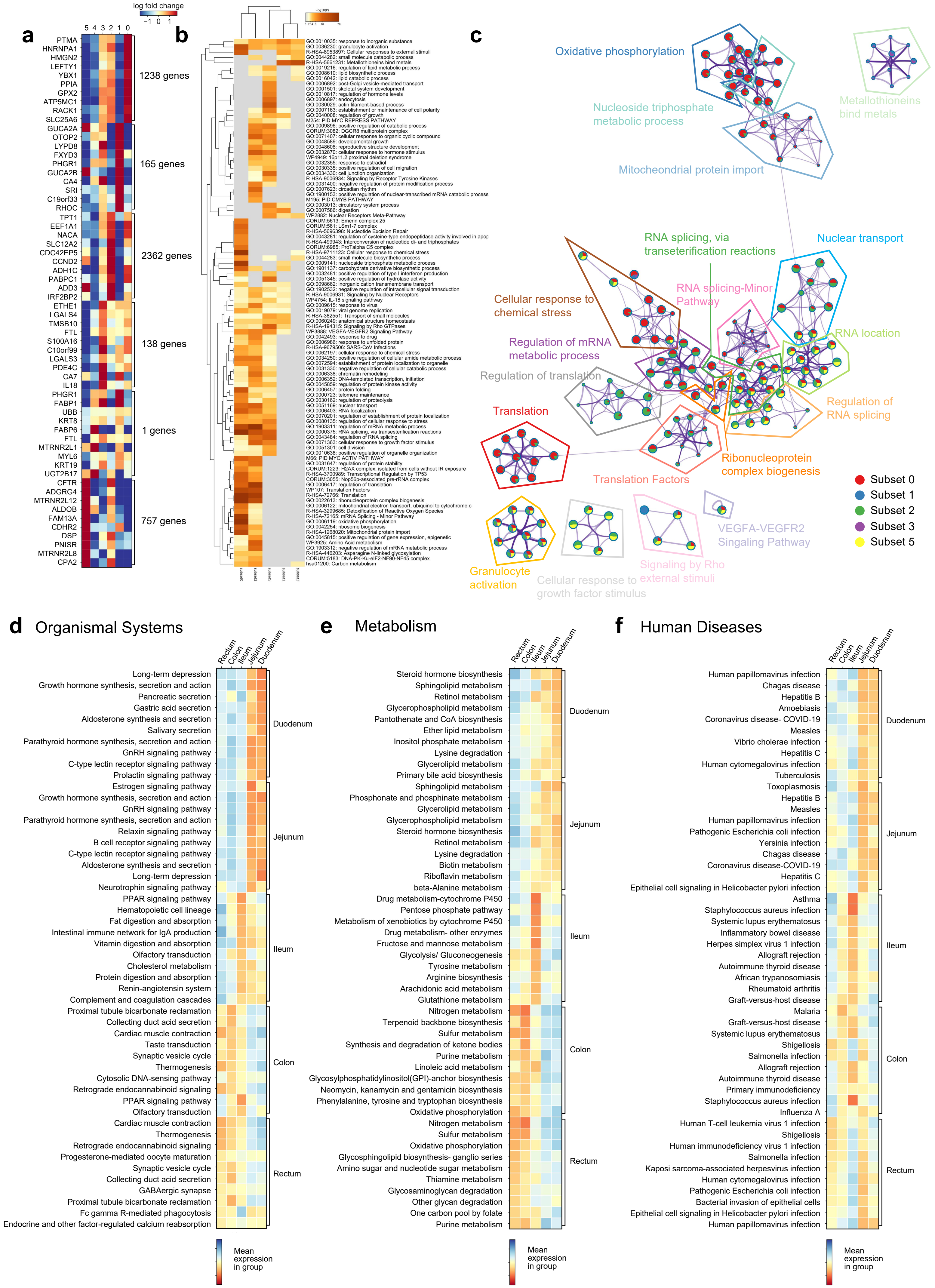

### Extended Data Fig. 9

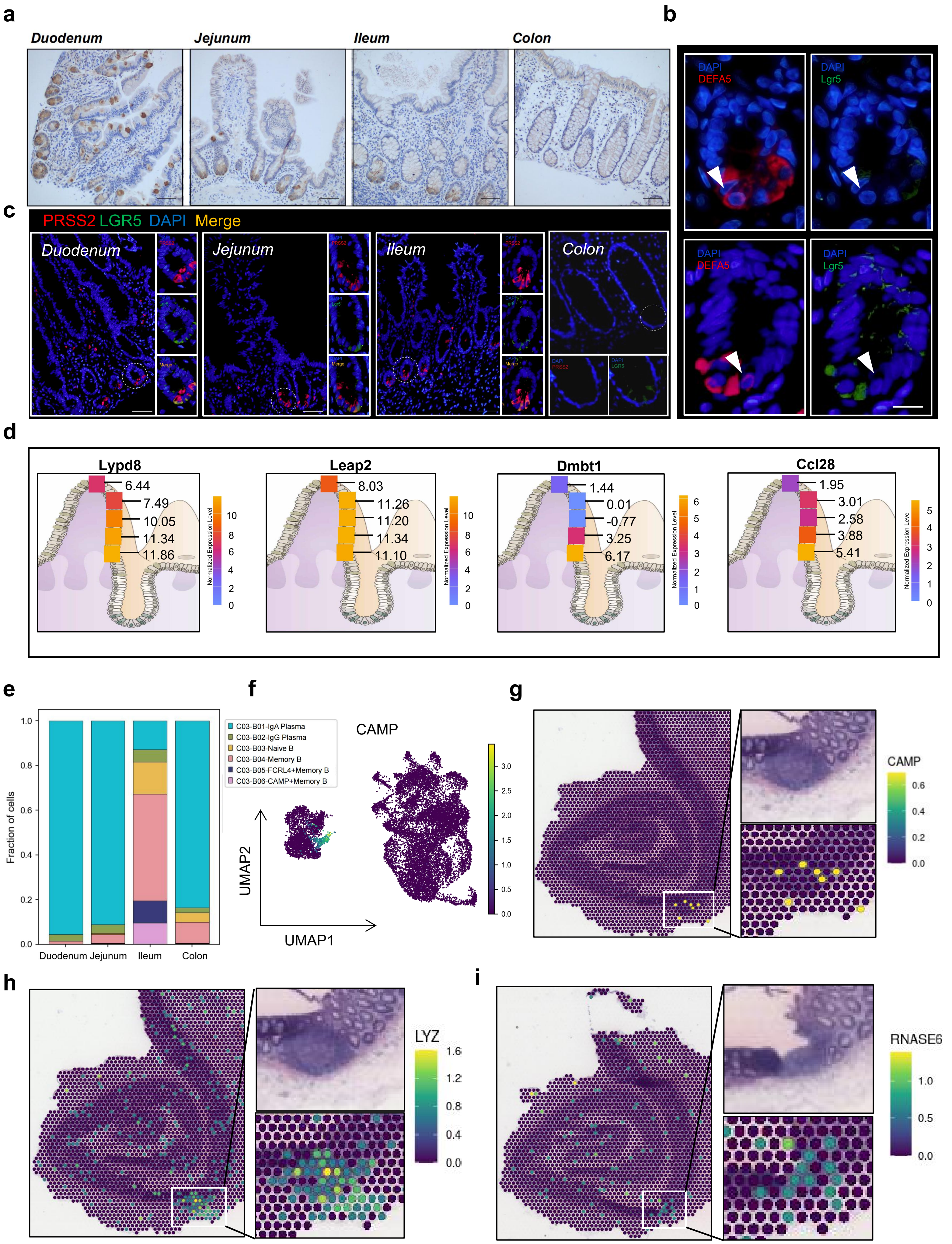

### Extended Data Fig. 10

a

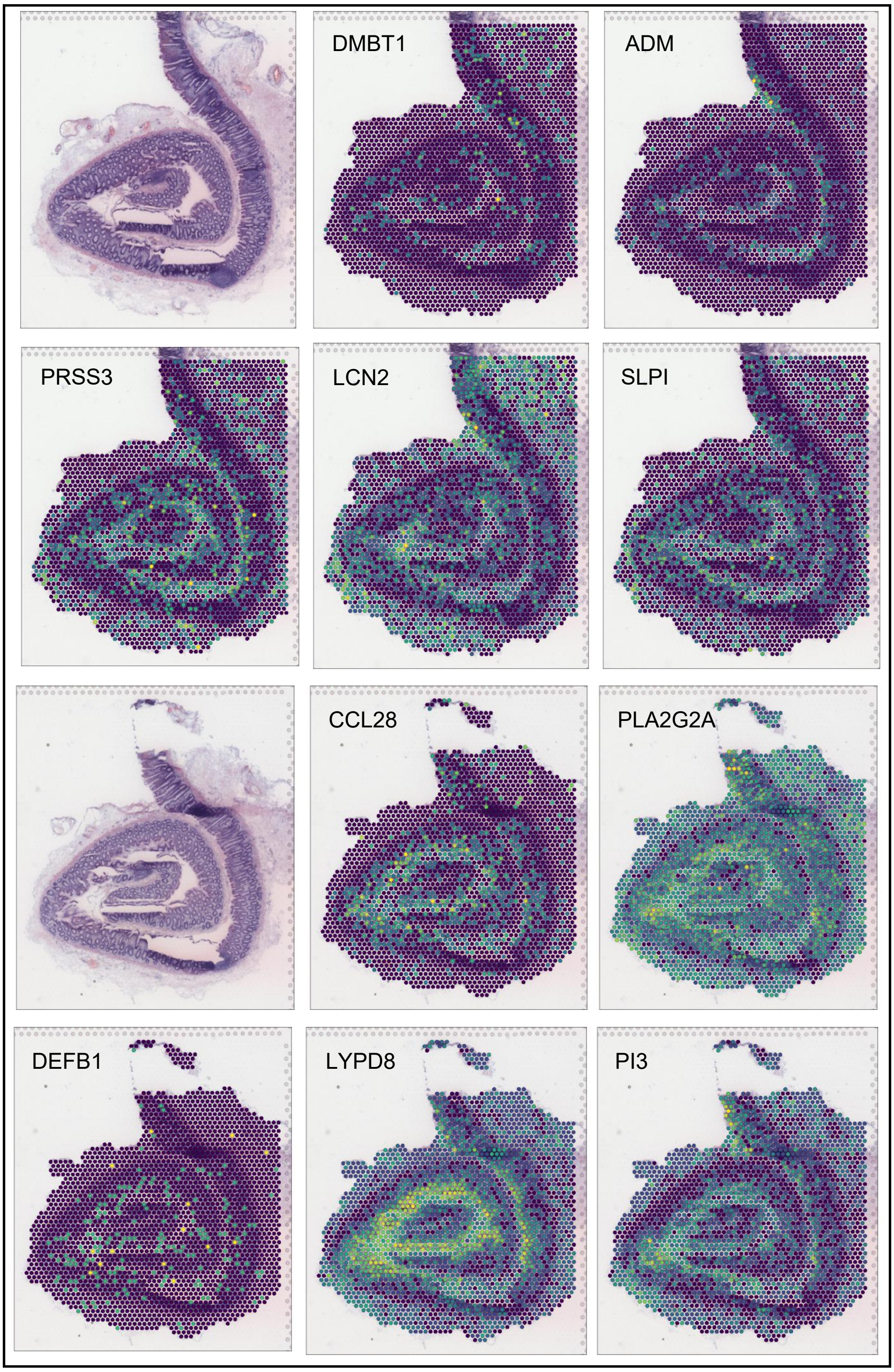


c

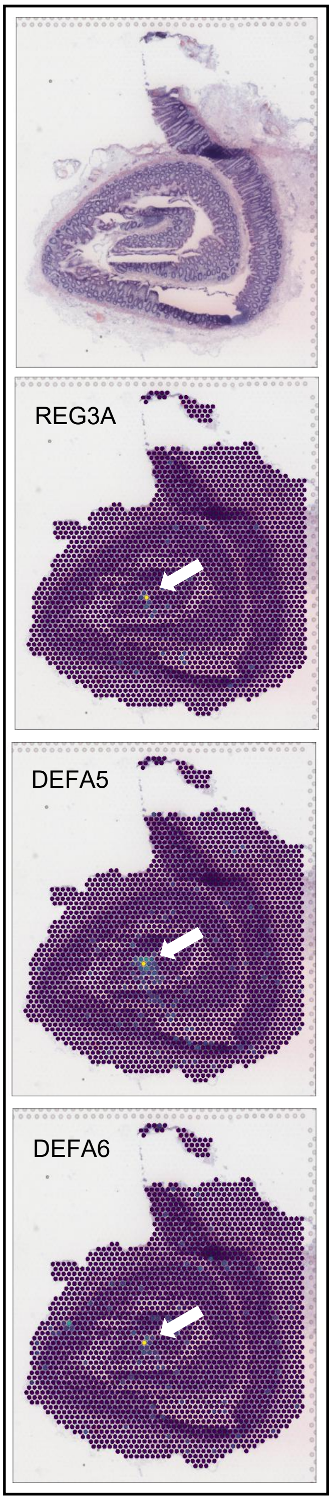


b

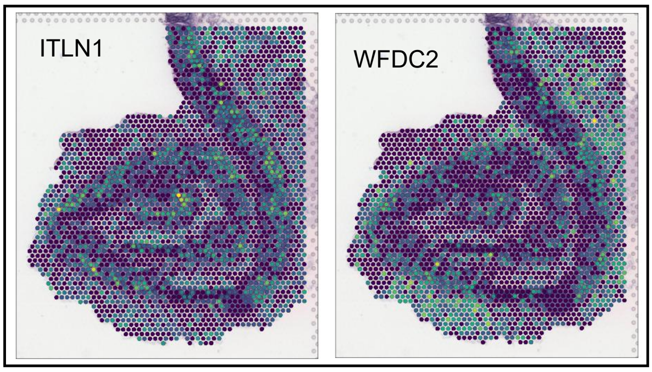

### Extended Data Fig. 11

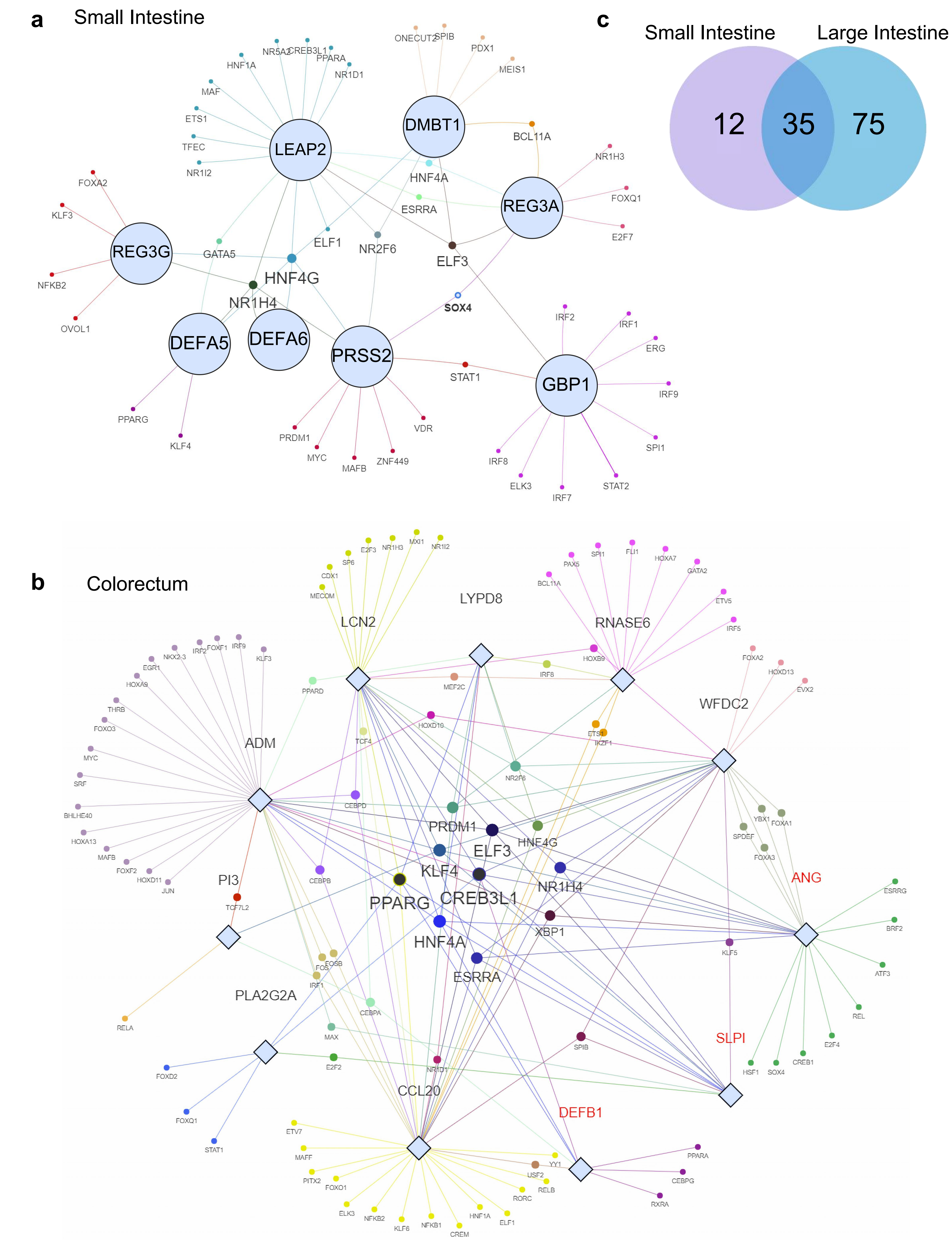

### Extended Data Fig. 12

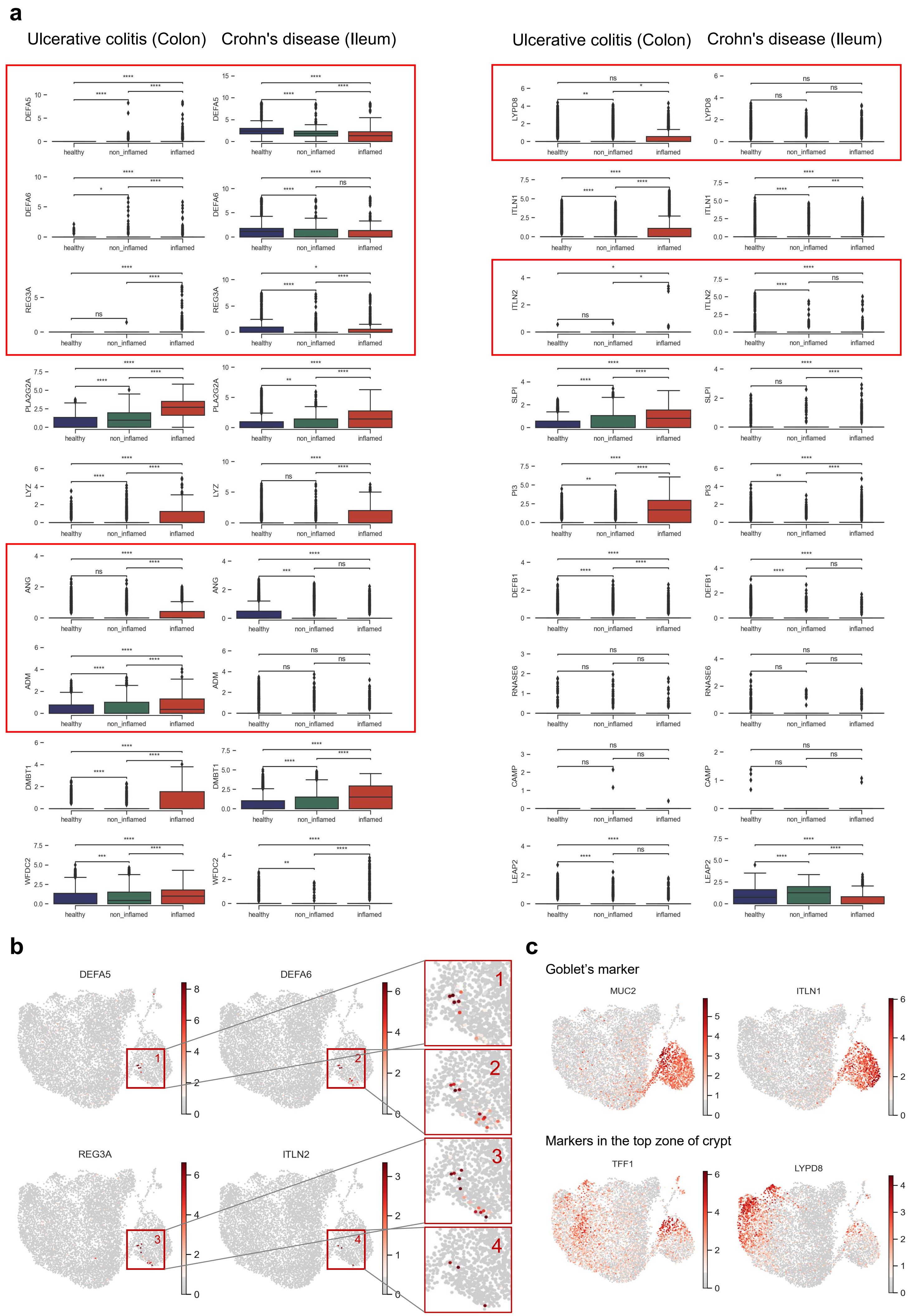

### Extended Data Fig. 13

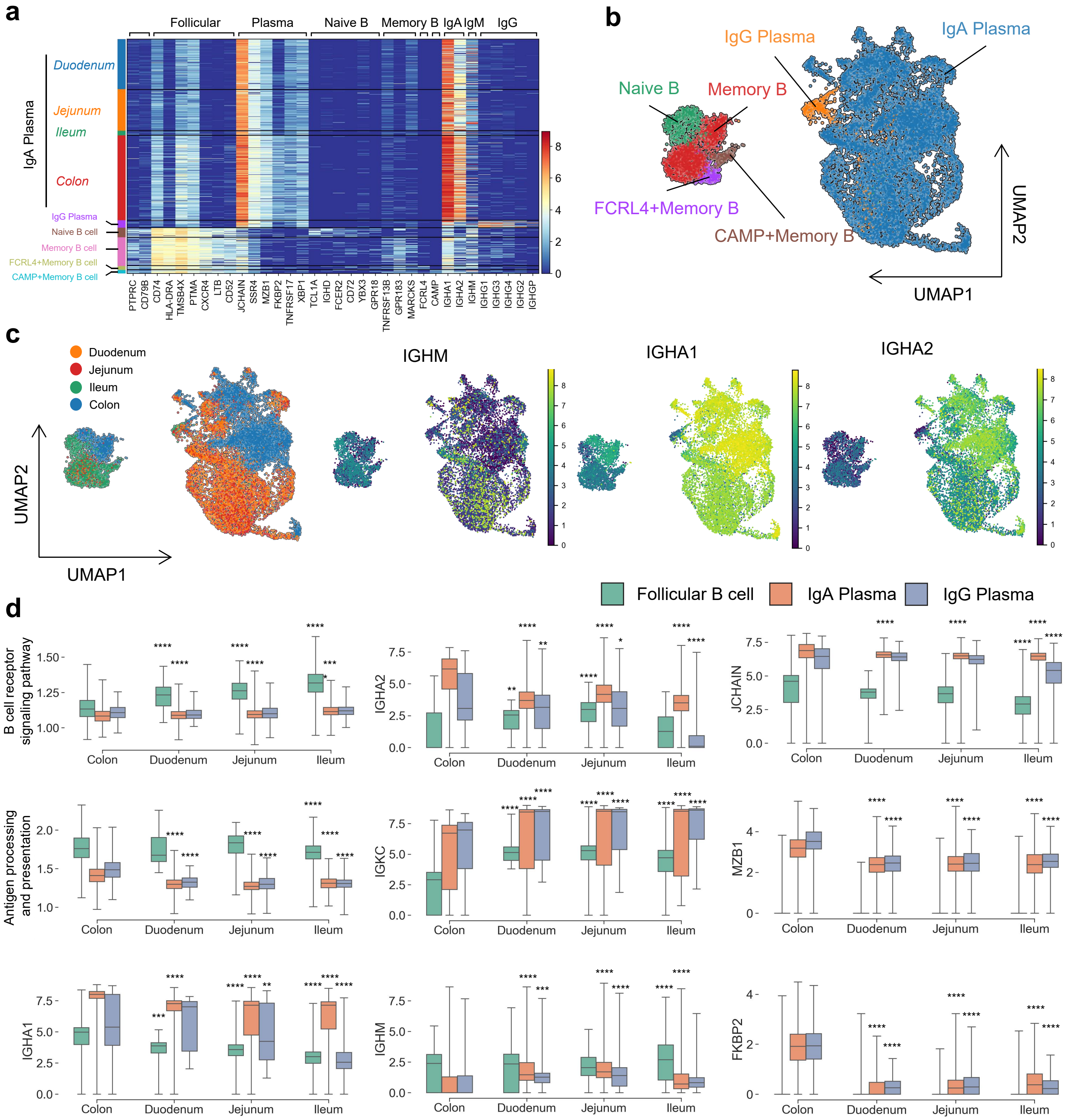

### Extended Data Fig. 14

**a**

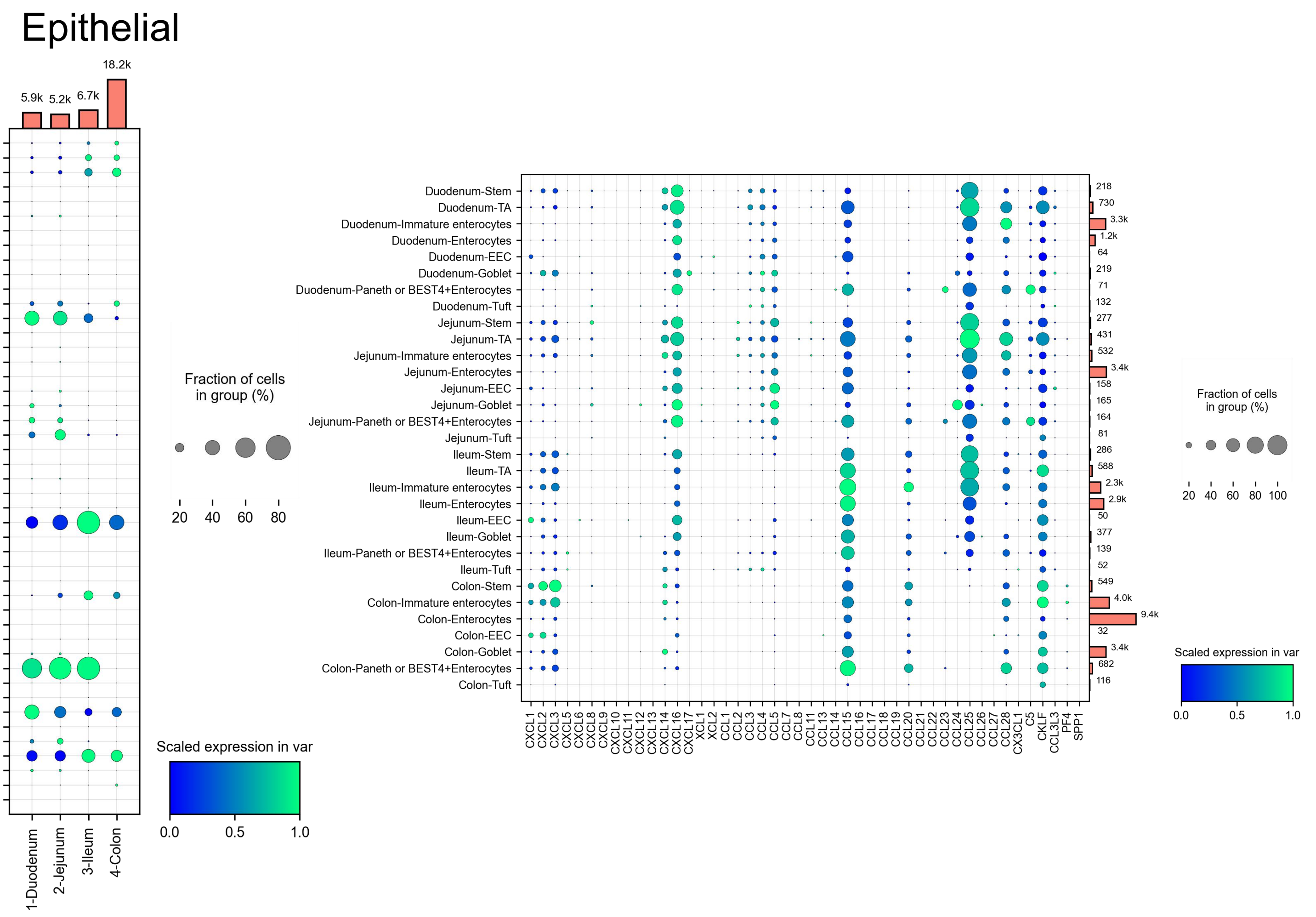


**b**

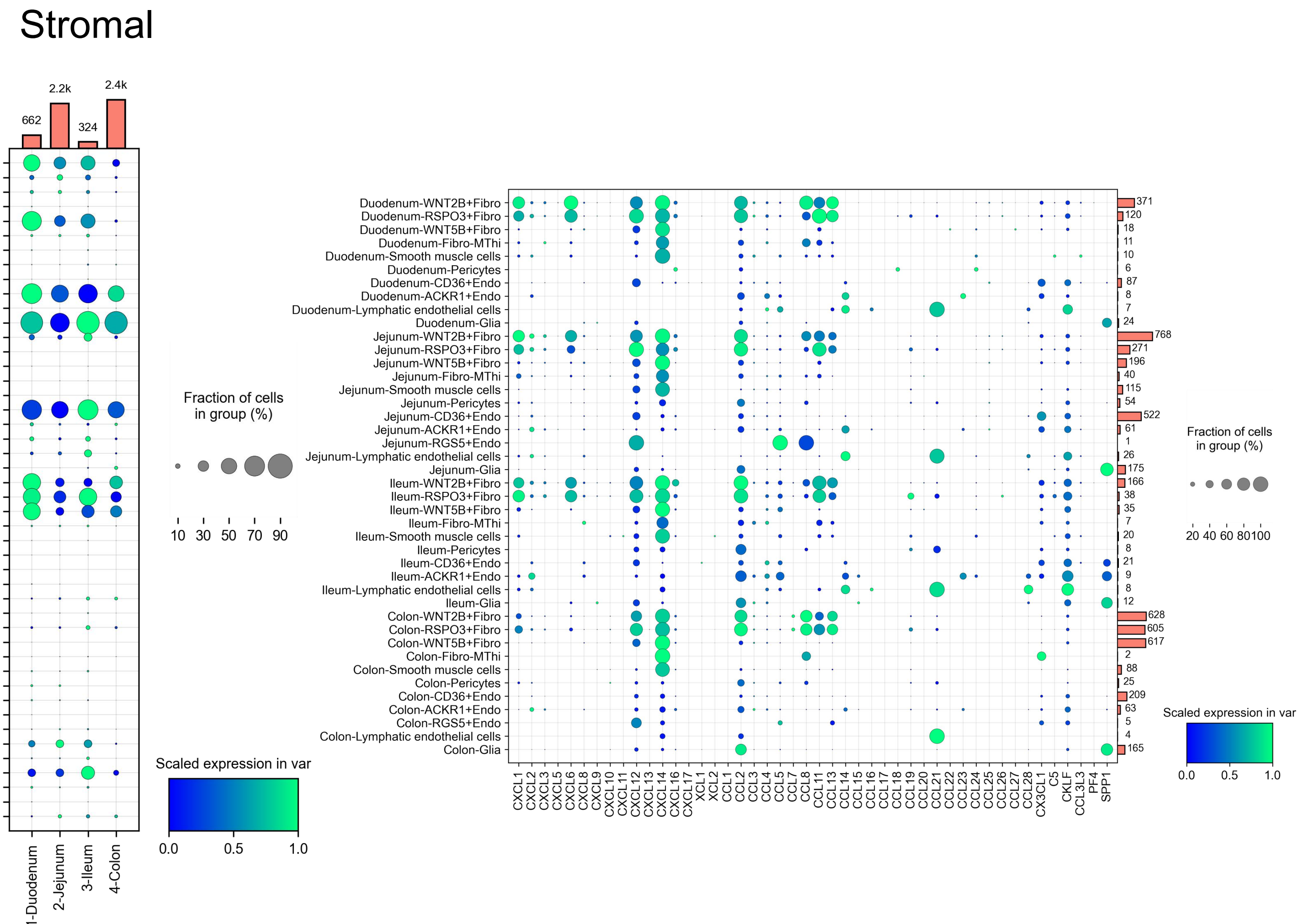

### Extended Data Fig. 15

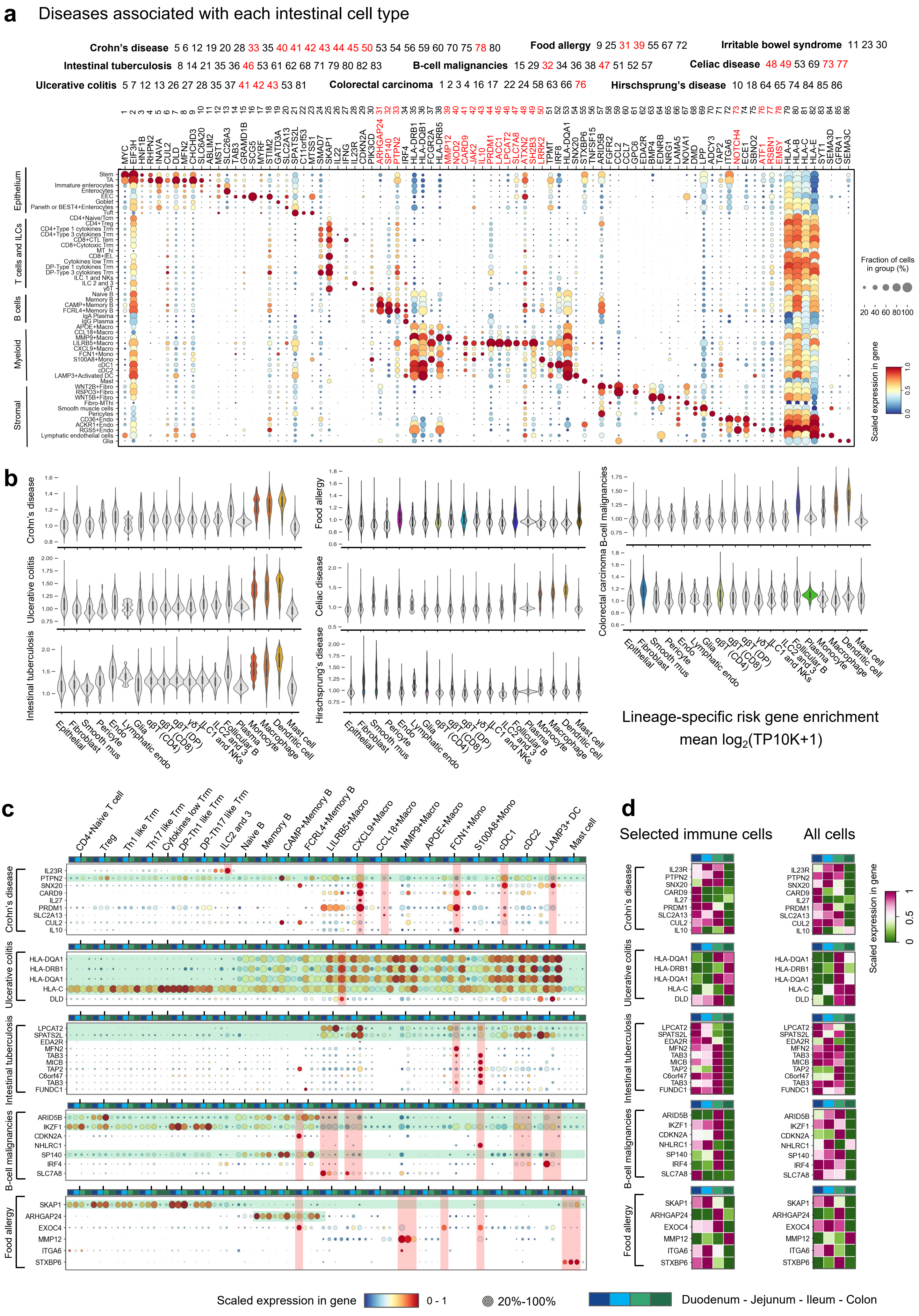
